# The Neonatal Gyrencephalic Cortex Maintains Regionally Distinct Streams of Neuroblasts

**DOI:** 10.1101/2023.06.06.543768

**Authors:** JaeYeon Kim, Kadellyn Sandoval, Aunoy Poddar, Julia Chu, Emma Horton, Di Cui, Keira Nakamura, Theresa Bartels, Christian Wood, David H. Rowitch, Hosung Kim, Chet C. Sherwood, Boris W Kramer, Angela C. Roberts, Pablo J. Ross, Duan Xu, Nicola J. Robertson, Peng Ji, Elizabeth A. Maga, Mercedes F. Paredes

## Abstract

Neurodevelopmental mechanisms have evolved to support the formation of diverse brain structures, such as in humans, during the perinatal period. Here, we demonstrate that neonatal gyrencephalic brains harbor an expanded subventricular zone, termed the Arc, defined by tiered arrangement of doublecortin (DCX)-expressing neuroblasts and vascular enrichment at the ventricular wall. The Arc is the origin of dorsal and ventral populations of migratory neuroblasts that target multiple regions involved in higher cognitive functions. Arc-derived migratory streams, primarily from the caudal ganglionic eminence, are composed of diverse neuronal subtypes with distinct spatial and migratory-receptor profiles. Our findings indicate the Arc is a structure present in phylogenetically divergent species that supports the expansion of postnatal neuronal migration, contributing to a protracted formation of cortical circuits in gyrencephalic brains.

**One-Sentence Summary:** The ventricular cytoarchitecture of gyrencephalic brains supports an ongoing supply of migratory neurons to the neonatal cortex.

## Main Text

Neuronal migration is a critical process through which immature neurons reach their anatomical destinations in developing brains (*1, 2*). This developmental stage is most robust in the prenatal period, however, there are restricted areas that maintain neuronal migration into the postnatal period. One example is the subventricular zone (SVZ), a neurogenic region that resides in the cortical ventricular wall and harbors young neuroblasts expressing doublecortin (DCX), a microtubule-associated protein fundamental to migration (*3-5*). The human SVZ has evolved into a complex organization, called the Arc, with DCX+ cells along the ventricular wall (tier1), dispersed away (tier2), around blood vessels (tier3), and as clusters oriented towards the pia in the developing white matter (tier4). Arc-derived neurons, primarily interneurons, target regions of the frontal cortex and the cingulate gyrus and disappear several months after birth (*6*). This analysis was restricted to the anterior ventricle in the human brain and whether there are additional migratory pathways remains unknown. Comparative studies on the mammalian SVZ suggest that this region has expanded within certain phylogenetic lineages of species. Neuroblasts in the rabbit brain, for example, organize into distinct clusters within the white matter closest to the SVZ (*7*). The marmoset, a nonhuman primate with a small, lissencephalic brain, has a lateral extension from the postnatal SVZ that maintains DCX+ cells for several months after birth (*8*). A small population of neuroblasts was observed in the dorsal parenchyma of these brains (*7-9*). The piglet SVZ has more complicated structure features including a vascular substrate, suggesting more similarity to human SVZ (*10, 11*). Understanding the evolution of this structure and how it contributes to migratory populations in the postnatal cortex is fundamental to identifying the development of cortical regions important for higher cognitive functions and their potential vulnerabilities to perinatal changes occurring before and after birth.

Here, we demonstrate that Arc features are present in gyrencephalic brains and serve as a postnatal reservoir for young migratory neurons. Additionally, using the piglet brain as a model for postnatal cortical migration, we identified dorsal and ventral migratory streams of primarily GABAergic interneurons targeting areas of the frontal, piriform, and temporal cortex during the early postnatal period. Spatial transcriptomics revealed the heterogenous molecular composition of these streams with a predominant representation of neurons derived from the caudal ganglionic eminence, a neurogenic niche in the ventral developing brain. Lastly, postnatal streams with distinct cortical destinations could be distinguished by their neuronal subtype composition, which showed differential expression of migration-related receptors. Our findings reveal robust neuronal migration into multiple cortical targets in neonatal gyrencephalic brain from diverse phylogenetic groups, serving as a potential mechanism for regulating function and plasticity across higher cognitive regions in the early postnatal period.

### A ventricular niche of migratory neurons in the developing gyrencephalic brains

We investigated the conservation of the Arc structure and DCX+ population at the ventricular wall in human, pig, marmoset, and mouse brains. Magnetic resonance imaging (MRI) classified the neonatal human and pig (piglet) brains as gyrencephalic, having cortical surface infoldings, and the marmoset and mouse brains as lissencephalic, with a smooth surface, even at adult ages (Fig. 1A, fig. S1A, and table S1) (*12, 13*). Serial section analysis of Nissl-stained coronal slices from the neonatal human and piglet ventricular wall showed a dense-cellular region, corresponding to the SVZ, that extended ventrally along the developing striatum; this was observed in the postnatal (P) 0 marmoset and mouse brains. The human and piglet SVZ also expanded from the dorsolateral wall with cellular extensions into the overlying white matter region. (Fig. 1, B, and C). The SVZ in macaques and chimpanzees, non-human primates, harbored a similar dorsal extension (fig. S1B). Based on their phylogenetic relatedness (*14*), it can be inferred that the gyrencephalic cortical surfaces of humans and pigs evolved independently. A correlation analysis comparing the Arc area and Gyrification Index (GI), a measure of the level of cortical folding, confirmed that the Arc area was significantly associated with GI (Fig. 1D, fig. S1C).

**Fig. 1.**
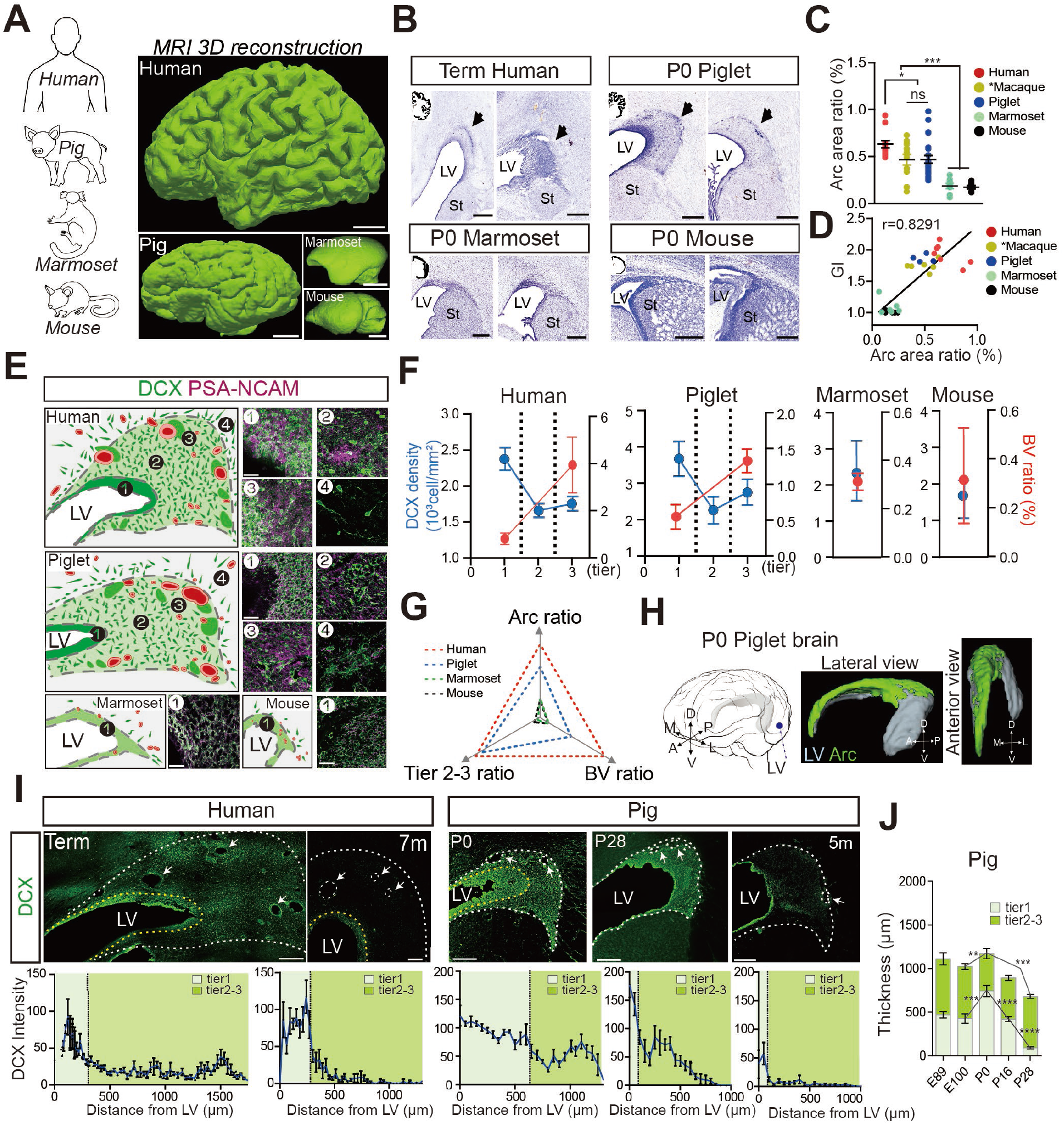
Arc structure is preserved in developing gyrencephalic brains. (**A**) MRI three-dimensional (3D) reconstruction of the postnatal human, pig, marmoset, and mouse brains. Two gyrencephalic brains (human at birth (term human) and P0 aged piglet) and two lissencephalic brains (marmoset at 4.5 years of age and mouse at birth). Scale bar, 1cm (human, pig, and marmoset); 1mm (mouse). (**B**) Nissl-stained serial sections across species taken at birth. Term humans and P0 piglets have a cell-dense region extending dorsally from the lateral ventricle (black arrows) that is absent in the P0 marmoset and P0 mouse. Scale bar, 500µm (human, piglet, and marmoset), 100µm (mouse); LV, lateral ventricle; St: striatum. (**C**) Quantification of Arc area relative to total brain area (Arc area ratio, %) by Nissl staining at birth. Macaque dataset (marked as *) from the public open source (NIH Blueprint NHP Atlas). Data means ± SEM (*, p < 0.05; **, p < 0.01; ***, p < 0.001 by paired t test). (**D**) Correlation (Pearson’s r) between Gyrification Index (GI) with the Arc area ratio across species (r=0.8291, p-value = <0.0001****). (**E**) Arrangement of DCX+ cells (in green) at the ventricular wall across species. Insets 1-4 (right of schematic) show confocal images of DCX+ enriched regions. Blood vessels are shown in red; dark green clusters correspond to DCX+ cellular densities. Scale bar, 30µm. (**F**) Quantification of DCX+ cells at the ventricular wall (blue line). Quantification of blood vessel (BV) areas, labeled by a-SMA, relative to total Arc area (BV ratio, %, red line) across species. In the human and piglet brains, DCX density and BV ratio are analyzed in each tier. In marmoset and mouse brains, they are analyzed in the ventricular wall. Data means ± SEM. (**G**) Triangle plot showing a comparison of core Arc features across species. Tier 2-3 ratio is calculated as the percentage of tier 2 and 3 areas of total Arc area. All values are normalized to humans (raw value and MANOVA analysis in sTable2). Red dotted line (human), blue dotted line (piglet), green dotted line (marmoset), and black dotted line (mouse). Core Arc features observed in the human are similar in the piglet (*F* (3,4) = 3.98, MANOVA *p*-value = 0.11), but significantly different from the marmoset and mouse (*F* (3,6) = 393.82, *p*-value = 6.21×10^-7***^; *F* (3,5) = 94.20, *p*-value = 8.07×10^-5***^; MANOVA analysis, respectively). (**H**) Schematic of the P0 piglet brain. The LV of the left hemisphere is highlighted in gray (left panel). 3D reconstruction of MRI images combined with immunohistochemistry for DCX and a-SMA (middle and right panel). The Arc (in green) extends in the rostral-caudal axis along the LV (in gray). (**I**) Top: Confocal image of DCX+ expression (green) in a coronal section of human and piglet Arc across early postnatal stages. The yellow and white dashed lines highlight the boundary between tiers 1 and 2 and tiers 3 and 4, respectively; arrows indicate blood vessels. Scale bar, 500µm. Bottom: Quantification of DCX expression pixel intensity across tiers. Tier 1 is separated from tiers 2-3 based on DAPI intensity (black dashed line). Data means ± SEM. (**J**). Quantification of the thickness of tiers from LV across ages of the pig. Data means ± SEM.

We previously showed that the human Arc contains migratory neurons that express DCX and PSA-NCAM (polysialylated neural cell adhesion molecule) and assemble in spatially distinct collections (*6*). In the piglet brain, DCX+ cells were grouped according to their distance from the ventricular edge, as was observed in the human Arc. In contrast, the majority of DCX+PSA-NCAM+ cells in the marmoset and mouse brain were arranged in a tangential fashion along the ventricular wall, indicative of solely having a tier1-like structure (Fig. 1E). Furthermore, the vascular area at the ventricular wall, a feature of Arc tier 3, was significantly reduced in P0 marmoset and mouse brains compared to neonatal humans and piglets (Fig.1F and fig. S1D). Multivariate analysis of variation (MANOVA) of core Arc features confirmed that the neonatal human brain was comparable with the piglet brain (*p*-value = 0.11), and significantly different from P0 marmosets and mice (*p*-value = 6.21×10^-7***^, *p*-value = 8.07×10^-5***^, respectively) (Fig. 1G and table S2).

Given the similarity between the neonatal human and piglet Arc region, we performed a comprehensive analysis of the piglet brain to understand the developmental dynamics of the postnatal Arc. 3D reconstruction using combined neuroimaging and histology showed that the P0 piglet Arc encompassed the entire body of the lateral ventricle (Fig 1H). This extended the previous study that was only able to capture anterior features of the human Arc (*6*). The tiers within the piglet Arc clearly appeared at P0 (fig. S2, A to C). The Arc area remained constant until P16 and dropped by 38% across the anterior-posterior axis at P28 (fig. S2D). This temporal pattern was mirrored by DCX intensity across Arc tiers and the tiers thickness (Fig 1, I and J, and fig. S2E). There was also a significant reduction in Arc volume of 40% at P28 (fig. S2F). At 5 months, DCX expression was rare in the ventricular region that had contained the piglet Arc, most notably in tier 2-3. The loss of Arc tiers in the piglet brain closely mimics the changes in the human Arc where DCX+ cells were distributed across tiers at birth and disappeared by 7 months, most dramatically within tiers 2-3 (Fig. 1I). Lastly, the density of glial and proliferative populations within the tiers was 25-fold less compared to the total DCX+ cell density, highlighting the Arc as containing mainly young migratory neurons (fig. S2G). Our data supports that the expanded ventricular niche with young migratory neurons termed the Arc, is present across developing gyrencephalic brains. The developmental comparison between the neonatal human and piglet brains demonstrated that the Arc is a transient structure in both species (fig. S2H).

### The Arc is a postnatal reservoir of diverse young GABAergic interneurons

GABAergic interneurons are essential for maintaining the balance of excitation and inhibition in the brain, and their maturation is involved in the acquisition of cognitive function (*15, 16*). Previous work demonstrated that DCX+ cells within the human Arc express GABAergic markers, consistent with an interneuron identity (*6*). Therefore, we asked whether the DCX+ cells within the postnatal piglet Arc would also be composed of interneurons. We performed single molecule fluorescent in situ hybridization (smFISH) using RNA probes against GABAergic interneuron markers (*DLX2* [Distal-Less Homeobox 2], *GAD1*, and *GAD2* [glutamic acid decarboxylase 1 and 2]), combined with the co-detection of DCX protein, on early postnatal human, piglet, and marmoset ventricular regions (Fig. 2A). Neuroblasts in the human and piglet Arc showed similar inhibitory representation, with 79.3% of DCX+ cells expressing *GAD1* in the human and 85.5% in the piglet. This proportion was preserved across Arc tiers (Fig. 2, B and C, fig. S3A). In the marmoset ventricular wall, about 75% of DCX+ cells were *GAD1*+. The density of DCX+ and *GAD1*+ cells at the ventricular wall was three-fold lower in the marmoset compared to in the human and piglet, consistent with a less expansive SVZ in the marmoset brain (fig. S3B)

**Fig. 2.**
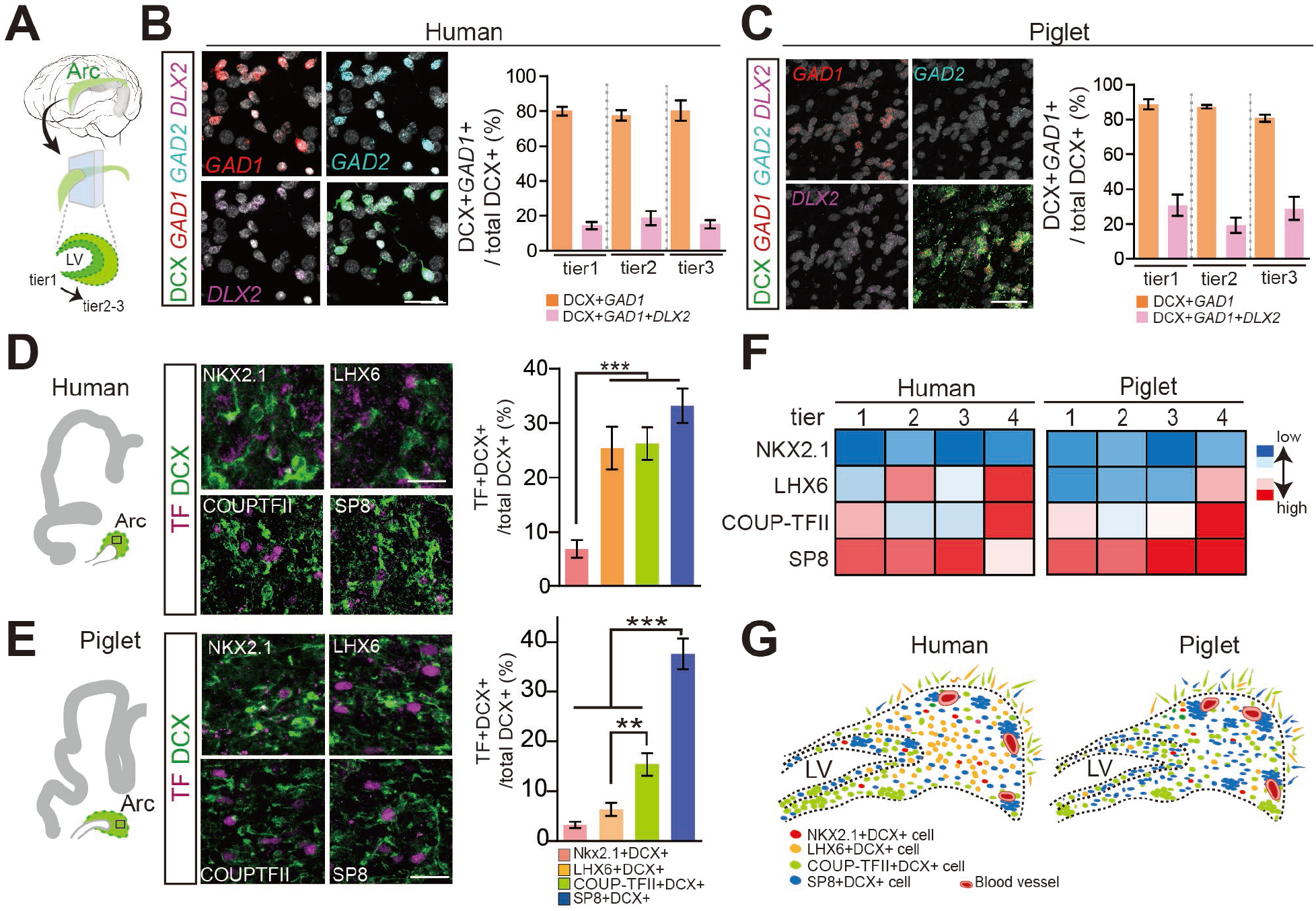
The Arc is a postnatal reservoir of diverse young GABAergic interneurons. (**A**) Schematic indicating areas within the Arc analyzed cellular composition. LV, lateral ventricle. (**B** and **C**). Confocal microscopy of smFISH for RNA expression of GABAergic interneuron markers (*GAD1, GAD2, DLX2*) in the neonatal human (B) and piglet brain (C). DCX protein expression was co-detected. Scale bar, 30µm. Right: Quantification of DCX+*GAD1*+ cells of all DCX+ cells across tiers. The majority of DCX+ cells in the neonatal human and piglet Arc are *GAD1*+. Data means ± SEM. (**D** and **E**) Left: Subpopulations of DCX+ cells in the neonatal human and piglet Arc express different transcription factors (TF) enriched in ventral telencephalic origins, immunostained with NKX2.1, LHX6, COUP-TFII, and SP8 associated with the MGE or CGE. Scale bar, 30µm. Right: Quantification of DCX+ cells expressing the selected TFs. Data means ± SEM of counts performed on three individual cases (n=3). (**F**) Heatmap of distribution of DCX+ cells expressing different TFs across tiers. The color gradient represents TF expression levels from high (red) to low (blue), as quantified from the counts performed on each species. (**G**) Schematic of the spatial distribution of molecularly distinct DCX+ cells in the neonatal human and piglet Arc.

GABAergic interneurons are a diverse neuronal subpopulation, primarily produced within ventral regions of the embryonic brain called the ganglionic eminence (GE), which can be divided into the medial, caudal, and lateral GE (MGE, CGE, and LGE, respectively) (*17-19*). Anatomical and transcriptomic features of the GE subregions are well conserved across species. The MGE located in medioventral GE is delineated by the expression of NKX2.1 (NK2 homeobox1) and LHX6 (LIM homeobox 6-positive); CGE located in caudoventral GE is marked by COUP-TFII (COUP transcription factor 2, also known as NR2F2 [nuclear receptor subfamily 2, group F, member 2]) expression (*20-24*). The subpopulation of dorsal LGE (dLGE)/CGE-derived neocortical interneurons express the SP8 (the zinc finger transcription factor) (*25, 26*). The piglet brain at embryonic day (E) 62, corresponding to the second trimester, maintained spatially restricted expression of GE molecular markers with NKX2.1, LHX6, COUP-TFII, and SP8 along the ventricular wall (fig. S4) (*27*). To identify the types of GABAergic interneurons and their distribution within the Arc, we analyzed the expression of GE-associated transcription factors (TFs) in DCX+ cells in the human and piglet ventricular wall. In the neonatal human Arc, about 6.9% of DCX+ cells were NKX2.1 and 25.4% were LHX6+. COUP-TFII and SP8 were expressed in 26.2% and 33.1% of DCX+ cells, respectively (Fig. 2D). In comparison, 3.2% of DCX+ cells were NKX2.1 and 6.3% were LHX6+ in the piglet Arc. COUP-TFII and SP8 were expressed in 15.3% and 37.6% of DCX+ cells, respectively (Fig. 2E and fig. S5A). We next asked how the composition of GE-derived DCX+ cells differed across Arc tiers. SP8+DCX+ cells were the most abundant subpopulation and present throughout the four tiers in both humans and piglets (Fig. 2F). In the human Arc, the next most abundant DCX+ subpopulations, LHX6+DCX+ and COUP-TFII+DCX+ cells, were primarily in tier 4 but also present in more proximal layers; tier 2 and tier 1, respectively. COUP-TFII+DCX+ cells in the piglet Arc were highly enriched in tier 4, as in the human. LHX6+DCX+ cells demonstrated the largest comparative difference with overall abundance being less in the piglet Arc and distribution mainly in tier 4. NKX2.1+ cells were the least abundant DCX+ subpopulation (fig. S5B). To further understand the spatial distribution of GABAergic interneuron subtypes in the P0 piglet Arc, we analyzed additional GE-associated molecular markers. DCX+ cells that express SCGN (expressed in CGE-derived cells) and PAX6 (expressed in pallial/dLGE-derived cells) were the most represented across Arc. PROX1+ (associated with the CGE), SCGN+, and PAX6+ DCX+ cells were populous in tier1. LGE-derived DCX+ cells that expressed FOXP2, GSH2, TSHZ1, and MEIS2, were a relatively small population and localized to tier 3 (fig. S5, C to G). These results indicate that the tiered Arc organization supports a large reservoir of diverse young GABAergic interneurons in the early postnatal gyrencephalic human and piglet brains (Fig. 2G).

### The Arc gives rise to expansive cortical streams of migratory neurons

We hypothesized that the Arc structure gives rise to multiple migratory populations in the postnatal cortex of gyrencephalic animals. Therefore, we mapped the distribution of DCX+ cells in the P0 piglet brain. Serial section analysis along the sagittal plane revealed that a thick band of DCX+PSA-NCAM+ cells around the body of the ventricle gives rise to distinct extensions along the anterior-posterior axis (fig. S6A). Two DCX+ streams emerged from the anterior olfactory ventricle (OV) at the most anterior level: the rostral migratory stream (RMS) that could be followed into the olfactory bulb (OB) and a thin medial stream that connected to the ventromedial region of the prefrontal cortex (PFC). More dorsally, migratory streams from the Arc targeted positions in the frontal cortex (Fig. 3A). These anterior migratory streams were analogous to those observed in the neonatal human brains (*6, 28-30*).

**Fig. 3.**
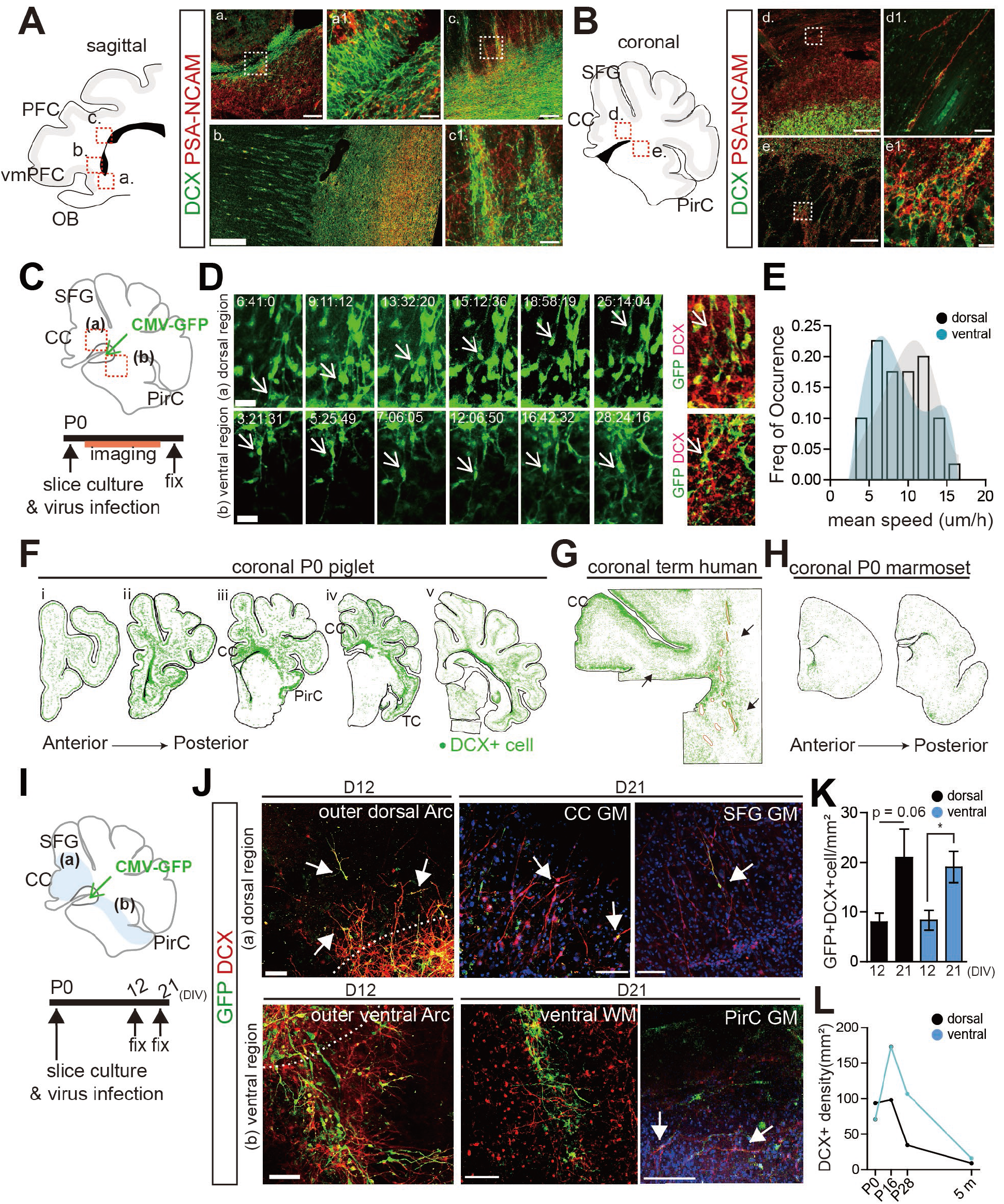
The Arc provides dorsal and ventral cortical streams of migratory neurons. (**A**) Left: schematic of sagittal section of the P0 piglet brain. Right: Migratory neurons expressing DCX (green) and PSA-NCAM (red) located in the RMS (a) targeting the OB, the streams targeting the vmPFC (b), and the streams from the Arc to the PFC (c). Scale bar, 100µm (a) and (c), 200µm (b), 20µm (a1) and (c1). (**B**) Left: schematic of a coronal section of the P0 piglet brain. Right: Typical appearance of the DCX+PSA-NCAM+ neurons as individual neurons in the dorsal Arc (d) and as cell clumps in the ventral Arc (e). Scale bar, 100µm (d) and (e), 20µm (d1) and (e1). (**C**) Experimental design for time-lapse imaging of the P0 piglet brain. Boxed areas (a) and (b), adjacent to viral injection sites (Ad-CMV-GFP), are further illustrated in (D). (**D**) Left: sequential images of time-lapse confocal microscopy showing GFP+ cells in the dorsal region (a) and the ventral region (b) of the piglet Arc. White arrows highlight GFP+ cells near the Arc. Right: Confocal image of GFP+ cells after 72 hours of imaging, showing GFP+ cells co-expressing DCX. Scale bar, 10µm. (**E**) Distribution of mean migratory speeds measured for the two populations of GFP+ cells from the dorsal region (n=21 cells; gray) and ventral region (n=20 cells, blue). (**F**) Mapping of DCX+ cells (green) in the P0 piglet brain. DCX+ cells from the Arc target the CC, PirC, SFG, and TC. (**G and H**) Mapping of DCX+ cells in term human coronal sections (G), at a plane that corresponds to (iv) in (F), as well as in the serial coronal sections of a P0 marmoset brain (H). The cortical streams from the Arc are shared by the neonatal human and P0 piglet brains, but not in the P0 marmoset brain. (**I**) Experimental design of extended time-lapse imaging performed on P0 pig brain slices. The number of GFP+ cells was quantified in a dorsal region, including the dorsal sides of the Arc and the CC (light blue (a)), and a ventral region, including the ventral sides of the Arc and the PirC (light blue (b)), at 12 and 21 DIV. (**J**) Confocal images of GFP+DCX+ cells in the dorsal (a) and ventral (b) regions at 12 and 21 DIV. White arrows indicate GFP+DCX+ cells with active migratory morphologies. Scale bar, 50µm. (**K**). Quantification of GFP+DCX+ cell density in the dorsal and ventral regions at 12 and 21 DIV, demonstrating GFP+DCX+ cells continue migrating at 21 DIV. Data means ± SEM of counts performed on three individual animals (n=3). (**L**) Quantification of DCX+ cell density in dorsal and ventral regions across different postnatal ages of the pig, equivalent to regions analyzed in (I). rostral migratory stream (RMS); olfactory bulb (OB); prefrontal cortex (PFC); ventromedial prefrontal PFC (vmPFC); cingulate cortex (CC); piriform cortex (PirC); superior frontal gyrus (SFG); temporal cortex (TC); gray matter (GM); white matter (WM); *days in vitro* (DIV).

In addition, the sagittal view demonstrated that neuroblasts reached the posterior end of the ventricular wall (fig. S6A), consistent with our 3D reconstruction (Fig 1I). These findings prompted us to examine the posterior piglet Arc and its contributions to other migratory streams. Using coronal serial section analysis (Fig. 3B), we observed separate and morphologically distinct streams along the outer edge of the Arc (tier 4). DCX+PSA-NCAM+cells at the dorsal side were radially oriented and mainly observed as individual cells with elongated morphologies. In contrast, DCX+ cells at the ventral edge of the Arc were organized as long clumps that stretched into lateral-ventral regions (Fig. 3B). These collections were often embedded in a matrix of brain lipid-binding protein (BLBP)+ cells (fig. S6B, and movies S1 and 2). DCX+ clumps were predominantly observed in the ventral stream with an average area of 11,934±300µm ^2^ and a total cell number of 75 or more, whereas those in the dorsal stream were small (average area of 4,000±1000µm ^2^; a total cell number of = 42 or more) and of low density (fig. S6, C to F).

To confirm that identified DCX+ populations were indeed migratory in the postnatal environment, we performed time-lapse confocal imaging on P0 piglet organotypic slice cultures. CMV-GFP adenovirus was microinjected into the Arc region of coronal cortical sections (Fig. 3C). Adenovirus-derived GFP was expressed within the Arc at 1 *day in vitro* (DIV) (fig. S7A). At 2 and 3 DIV, we observed GFP+ cells in both the dorsal and ventral subregions with a small, elongated nucleus and a leading process directed away from the Arc (Fig. 3D, movie S3 and 4). Post hoc immunostaining of these slices after time-lapse imaging showed that migratory GFP+ cells were DCX+ (Fig. 3D and fig. S7B). To characterize the physical migratory behaviors of the postnatally migrating cells, we tracked 41 cells from the dorsal and ventral outer Arc. The cells in the dorsal and ventral outer Arc did not differ in speeds, having an average of 8.82µm/h and 7.5µm/h, respectively (fig. S7C). Migratory profiles and frequency distribution of mean speed suggested a trend toward more heterogeneous behaviors in the ventral subpopulation (Fig. 3E and fig. S7D).

DCX+ mapping along the coronal plane showed that the dorsal and ventral Arc had unique cortical trajectories (Fig. 3F). The dorsal Arc gave rise to a DCX+ stream that connected with the cingulate cortex (CC) and superior frontal gyrus (SFG) (fig. S8A). DCX+ cells from the ventral Arc extended as chains of cell clumps that separated into individual cells within the piriform cortex (PirC) and temporal cortex (TC) (fig. S8B). Young neuroblasts in the neonatal human brain also had a ventral extension from the Arc as cell clumps (Fig. 3G and fig. S8C). In contrast, the P0 marmoset brain had few individual DCX+ cells with migratory morphology at ventral and dorsal SVZ regions (Fig. 3H and fig. S8D).

We tracked GFP+ cells over 21 DIV to confirm their cortical trajectories from the piglet Arc (Fig. 3I). GFP+DCX+ cells were seen dorsally and ventrally near the outer Arc by 12 DIV. GFP+DCX+ cells exited the dorsal Arc as individual cells and had radial morphology; GFP+DCX+ cells at the ventral Arc collectively migrated as cell clumps. At 21 DIV, GFP+DCX+cells were observed in CC and SFG gray matter (GM), ventral white matter (WM), and in PirC (Fig. 3J). Quantification of GFP+DCX+cells supported their influx into both dorsal and ventral regions between 12 and 21 DIV (Fig. 3K). Coronal section analysis at P16 and P28 showed that the ventral stream remained detectable while the dorsal stream was depleted; DCX+ cells within streams were absent by 5 months, indicating that populations from the dorsal and ventral Arc were transient (Fig. 3L and fig. S9). This data demonstrates that there are distinct early postnatal DCX+ streams from the Arc into several cortical areas in human and piglet brains (movie S5).

### Migratory neurons in the neonatal cortex are derived predominantly from the Caudal Ganglionic Eminence

We next profiled GE-associated TFs in DCX+ cells in the migratory streams from the piglet brain to determine the subtype composition within Arc-derived cortical streams (Fig. 4A). The dorsal streams into the CC contained a dominant population of LHX6+ and COUP-TFII+SP8+ DCX+ cells, whereas the ventral streams into the TC were most highly populated by COUP-TFII+SP8+ DCX+ cells. The streams into the PirC were equally populated by the above subtypes (Fig. 4, B, and C). SP8+DCX+ cells that did not express COUP-TFII were abundant in the RMS, as well as in OB, as reported in the mouse and human (*22*) (fig. 10A). COUP-TFII+SP8+ cells were the most abundant subtype in posterior cortical streams, at 43.5 % of the DCX+ population (fig. 10B). SP8+DCX+ and COUP-TFII+DCX+ cells, in addition to LHX6+ and NKX2.1+ cells, were rare (fig. 10B, Fig. 4, B and C).

**Fig. 4.**
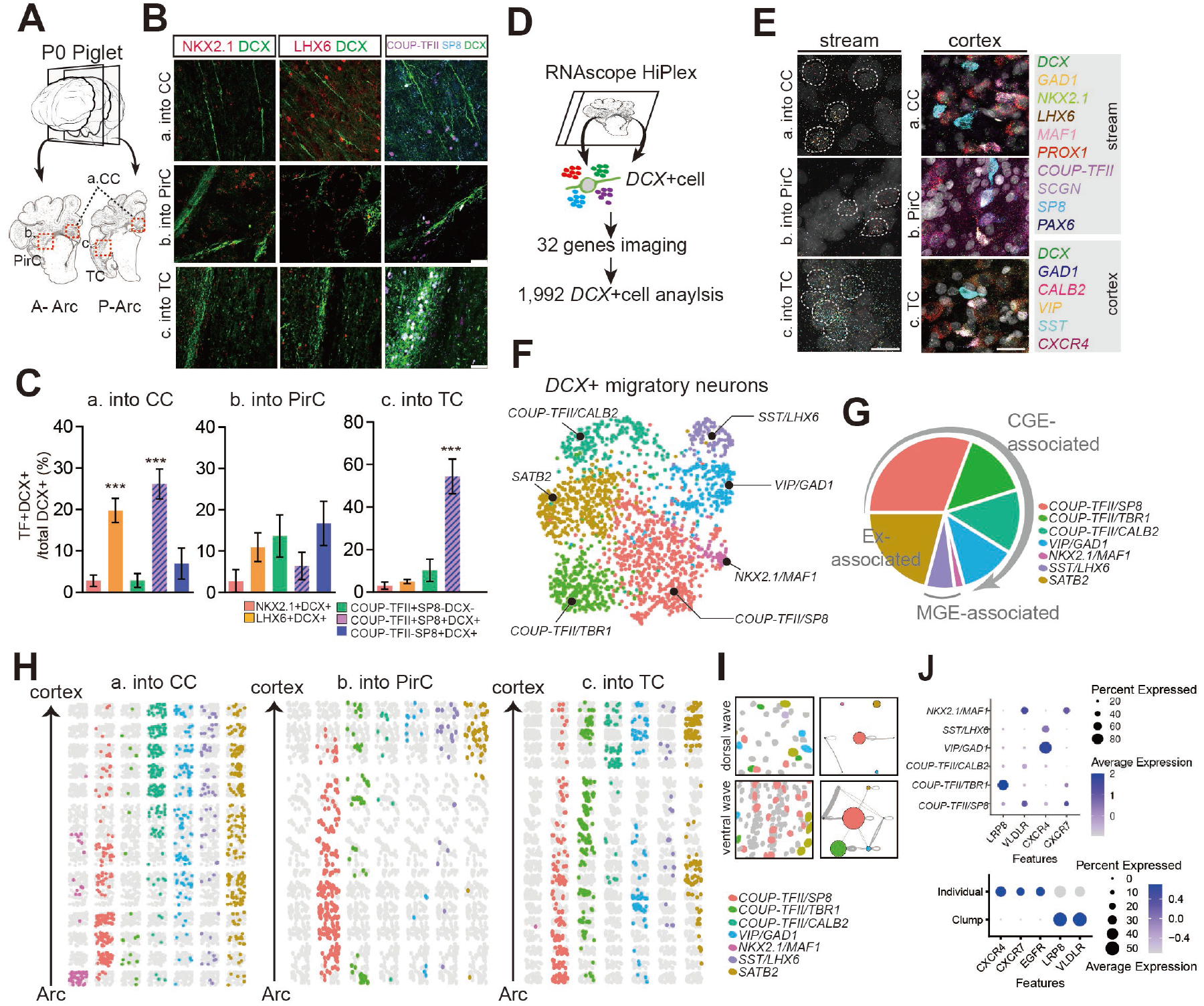
Migratory neurons in postnatal cortical streams predominantly express CGE-associated transcription factors. (**A**) Schematic indicating the postnatal migratory streams analyzed for profiling cellular subtypes in the P0 piglet brain: dorsal migratory streams into the anterior and posterior CC from the Arc; b: ventral streams into the PirC from the Arc; c: ventral streams into the TC from the Arc. (**B**) DCX+ subpopulations in each migratory stream express different transcription factors enriched in ventral telencephalic origins, including NKX2.1 and LHX6 (associated with the MGE) or COUP-TFII and SP8 (CGE). Scale bar, 50µm. (**C**) Quantification of DCX+ cells expressing selected TFs. Data means ± SEM of counts performed on three individual cases (n=3). (**D**). Schematic of showing the experimental design for P2 piglet spatial transcriptomics. (**E**). Left: smFISH confocal images of each stream showing examples of *DCX*+ cells expressing *GAD1, NKX2*.*1, LHX6, MAF1, PROX1, COUP-TFII, SCGN, SP8, and PAX6*, which are associated with GE. Right: smFISH confocal images of each cortical region showing examples of heterogeneous interneuron types expressing *GAD1, CALB2, VIP, SST, and CXCR4*. Scare bar, 25µm. (**F**) UMAP projections colored by cell identity. n = 1,992 *DCX+* cells. (**G**) Pie chart showing the proportion of clusters grouped by associated with ganglionic eminence region. Over 70% of cells express CGE-associated *COUP-TFII*, and 8.2% express MGE-associated *NKX2*.*1*, and *LHX6*. Others express *SATB2*, indicating excitatory neuron identity. (**H**) Topographic mapping of migratory neuron subtypes in each stream. From bottom to top shows the migratory stream from the Arc to their final cortical region, i.e., CC (left), PirC (center), and TC (right). (**I**) Nearest neighbor analysis. Gray line indicates a significant interaction between cell subtypes. The line thickness is correlated with the distance between the cell subtypes. Colored circles represent each cell subtype. A larger circle size indicates a higher number of cells within the cell subtype. The ventral streams show a higher interaction between cell subtypes. (**J**) Top: dot plot illustrating expression pattern of receptors for neuronal migration across subtype clusters. Botton: dot plot illustrating expression pattern of receptors across individually and clump migrating cells. cingulate cortex (CC); piriform cortex (PirC); temporal cortex (TC); anterior Arc (A-Arc); posterior Arc (P-Arc); transcription factor (TF); ganglionic eminence (GE); excitatory neuron (Ex).

To further define the molecular profiles of neuroblasts in the postnatal cortex, we performed multiplex smFISH on the P2 piglet cortex. We individually selected *DCX*+ cells as regions of interest (ROIs) and measured the ratio of puncta area expressed by different genes in the *DCX*+ cells to obtain the transcriptional profiles of 1,992 single *DCX*+ cells (Fig. 4, D, and E). Through unsupervised clustering of cellular transcriptional identities by uniform manifold approximation and projection (UMAP) dimensionality reduction, we defined 7 cell subclusters; *NKX2*.*1/MAF1, SST/LHX6, COUP-TFII/SP8, COUP-TFII/TBR1, COUP-TFII/CALB2*, and *VIP/GAD1*. A *SATB2* (excitatory neuron) cluster was also defined, which made up 22% of the *DCX*+ cells (Fig. 4F, fig. 11, A and B). Over 70 % of *DCX*+ cells expressed *COUP-TFII*, while 8.2 % expressed *NKX2*.*1* or *LHX6*, indicating the prominence of CGE-derived interneurons in the postnatal migratory population (Fig. 4G, fig. 11, C and D).

To examine the spatial distribution of the *DCX*+ subtype clusters, we performed topographic mapping (Fig. 4H). *COUP-TFII* was expressed by different *DCX*+ subpopulations in both migratory streams. The *COUP-TFII/TBR1* cluster was most abundant in the ventral stream into the TC and the *NKX2*.*1/MAF1* cluster was restricted to the dorsal stream into the CC. *COUP-TFII/CALB2, VIP/GAD1*, and *SST/LHX6* clusters, expressing markers of mature interneuron subtypes, were more abundant within the cortex, most notably in CC (fig. S11, E, and F). This suggests that postnatal migratory interneurons mature along their cortical trajectories. Nearest neighbor analysis showed that cells in the ventral stream had more interactions compared to those in the dorsal stream, consistent with ventral cells more frequently forming cell clumps. The *COUP-TFII/SP8* and *COUP-TFII/TBR1* clusters in the ventral stream, which had the highest interaction value, also had high self-interaction, indicating that they formed homogenous cell clumps (Fig. 4I). Lastly, we analyzed the spatial expression of receptors implicated in interneuron migration. *VLDLR* (very low-density lipoprotein receptor) and *LRP8* (low-density lipoprotein receptor-related protein 8; also known as apolipoprotein E receptor 2 [APOER2]) are essential receptors for Reelin, a key extracellular matrix protein for migration of interneurons (*31*). *VLDLR* was more expressed in *NKX2*.*1/MAF1* and *COUP-TFII/SP8* cluster, whereas *LRP8* was expressed by the *COUP-TFII/TBR1* cluster. *CXCR4* and *CXCR7* are known receptors for *CXCL12* (also known as stromal-derived factor 1 [SDF1]) expressed by MGE and CGE-derived migratory interneurons (*32*). *CXCR4* was expressed by both *SST/LHX6* and *VIP/GAD1* clusters, while *CXCR7* was the least expressed, observed in *NKX2*.*1/MAF1* and *COUP-TFII/SP8* clusters. Receptor expression was most distinct when cells were separated into those arranged individually, expressing *CXCR4* and *CXCR7*, and those arranged in cell clumps, expressing *LRP8* and *VLDLR* (Fig. 4J). Thus, chemokine-receptors differentially expressed in each migratory stream could be key spatial regulators, responding to mechanical cue for the postnatal integration of migratory interneurons from the Arc into their final cortical destinations.

## Discussion

We identified that an enlarged and tiered SVZ, termed the Arc, is present across the gyrencephalic brains. The neonatal human and piglet Arc share core features such as a layered arrangement of young neuronal populations, not observed in rodents and marmosets. Whether the occurrence of Arc is due to conserved mechanisms that are not fully expressed in smaller lissencephalic brains or has arisen through independent evolution in humans and piglets remains to examined. Extensive migration of predominantly CGE-derived neuroblasts from the Arc targeted regions involved in higher cognitive functions, including the frontal, piriform, and temporal cortex, in the neonatal piglet brain. These migratory routes were also observed in the infant human brain. Our findings support the Arc as a postnatal reservoir of diverse interneuron subpopulations that support the expansion of postnatal neuronal migration in larger, gyrencephalic animals.

Our data provide evidence that the development of CGE-derived interneurons, especially their postnatal migration, scales across species. Experiments on the 5HT3a-GFP mouse showed CGE-derived migratory neurons target several regions, including the dorsal cortex and striatum through P10 (*33, 34*); DCX+ neurons in the P20 ferret brain, a species that first develops gyri postnatally, express CGE-related markers, such as SP8 and SCGN in small migratory streams to PFC and the occipital lobe (*35*). Here, we found that over half of DCX+ cells within the piglet and human Arc express SP8 and COUP-TFII, consistent with previous reports in the piglet brain (*10*). Spatial transcriptomic profiling confirmed that the majority of DCX+ neurons in the cortical streams from the Arc were also CGE-derived. The expansion of late migratory CGE-derived neurons could contribute to a larger proportion of CGE-derived interneurons in the human brain (*21, 22, 36*). We also identified a subpopulation of CGE-derived neurons expressing *TBR1* enriched in the ventral migratory stream targeting the piriform and temporal cortex. The prenatal rodent brain contains a population of migratory cells at the cortico-striatal junction with a similar extension into the piriform cortex and amygdala (*37*). This collection termed the lateral cortical stream (LCS), supplies a population of cells that express TBR1, PAX6, and GSH2, supporting both excitatory and inhibitory identities (*38, 39*); the nature of TBR1+ cells from the CGE and the evolutionary relationship between the ventral stream from the Arc and the LCS remains to be elucidated. In comparison to CGE, the LGE contribution to the postnatal migratory streams is smaller. Expression of LGE-associated genes has been reported in neurons of the macaque cortex (*40*). These populations were present at late gestational ages and associated with LGE-derived populations from the early macaque that were similarly observed in the human Arc. Thus, the LGE contribution to cortical development could arise at earlier stages than the CGE.

How do young migratory neurons from the Arc target their final cortical destinations and how might this process be disrupted? Differential receptor expression might regulate postnatal migratory pathways. For example, *VLDLR* was primarily expressed in MGE- and CGE-associated clusters of the dorsal stream, whereas *LRP8* was in the *COUP-TFII/TBR1* cell cluster of the ventral stream. CXCR4 and CXCR7 were enriched in radially migratory DCX+ cells. Their divergent expression within migratory streams could coordinate how populations leave the Arc in unique migratory streams. The recruitment of Arc-derived neuroblasts both provides a mechanism to postnatally influence plasticity and has neuroprotective implications given that perinatal injuries, such as hypoxia-ischemia (*41*), could impact the survival and function of these late-migratory neurons, with long term cognitive outcomes.

## Funding

National Institutes of Health grant P01 NS083513 (M.F.P., D.H.R)

National Institutes of Health grant DP2 NS122550-01 (M.F.P.)

Roberta and Oscar Gregory Endowment in Stroke and Brain Research (M.F.P)

Chan Zuckerberg Initiative (M.F.P)

Weill Award for Junior Investigators in the Neurosciences Impacted COVID-19 Setbacks (JY.K.)

UCSF/UCB Schwab Dyslexia & Cognitive Diversity Center, 2023 Innovation Fund (JY.K.)

National Institutes of Health grant NS092988 (C.C.S.)

Medical Research Council UK MR M006743 1 (N.J.R)

## Author contributions

Conceptualization: K.S., M.F.P., JY. K., A.P.

Methodology: A.P., D.C., D.X.

Investigation: K.S., JY.K., A.P., J.C., E.H., K.N.

Visualization: JY.K., A.P., H. K

Funding acquisition: M.F.P., JY.K., C.C.S., N.J.R.

Project administration: E.A.M., M.F.P. Supervision: M.F.P.

Writing – original draft: JY.K., M.F.P.

Writing – review & editing: JY.K., A.P., T.B., C.W., D.H.R., H.K., C.C.S., B.W.K., A.C.R., P.R., D.X., N.J.R., P.J., E.A.M., M.F.P.

## Data and materials availability

All data are available in the main text or the supplementary materials.

## Supplementary Materials

Materials and Methods

Supplementary Text

Figs. S1 to S11

Tables S1 to S5

References(*42-48*)

Movies S1 to S5

