## Supplement for "The Neonatal Gyrencephalic Cortex Maintains Regionally Distinct Streams of Neuroblasts"

#### Materials and Methods

*Tissue collection*: Six de-identified human specimens were collected during autopsy, with a post-mortem interval of fewer than 48 hours (Table S3). Tissue was collected with previous patient consent in strict observance of the legal and institutional ethical regulations of the University of California San Francisco Committee on Human Research. Protocols were approved by the Human Gamete, Embryo, and Stem Cell Research Committee and the Committee on Human Research (institutional review board) at the University of California, San Francisco. Chimpanzee brains at birth (n=2) were provided by National Chimpanzee Brain Resource. Pig brains from embryonic day (E)62, E89, E100, postnatal day (P) 0-2, P16, P28, and 5 months of age (n=2-5) were collected at the Swine Teaching and Research Center at the University of California, Davis. All animal procedures conformed to the requirements of the Animal Welfare Act and were carried out under the Association of Assessment and Accreditation of Laboratory Animal Care International (AAALAC) approved conditions with protocols approved before implementation by the Institutional Animal Care and Use Committee (IACUC) at the University of California, Davis. Preterm and P0 marmoset brains (n=1-3) were collected immediately following euthanasia for welfare purposes at the University of Cambridge Marmoset Breeding Colony in accordance with the local Animal Welfare and Ethical Review Board. The P0-aged mice (C57/BL6) (n=3) were collected at the University of California, San Francisco (UCSF IACUC protocols AN192603-01) and were anesthetized on ice for 1-2 minutes. Preterm marmoset brains were fixed in 10% formalin in PBS for 1 day. All other brains were cut as a coronal or sagittal block and fixed in 4% paraformaldehyde (PFA) in PBS for 2 days (P0-aged mice for 1 day) at 4 °C. Cryopreservation was in 30 % sucrose in PBS at 4°C, until tissue specimens had sunk to the bottom of the vial. Tissue specimens were cut into coronal or sagittal blocks, and frozen in an optimal cutting temperature (OCT) compound. Blocks were cut at ~30 µm on a cryostat and mounted on glass slides for experimental analysis.

*MRI*: The images were acquired on a full body GE MR950 7T scanner using a 32-ch Nova Medical head coil with 3D fast spin echo sequence, isotropic 600-micron resolution, time to echo (TE) of ~120ms and TR of 2.5s, and 8 averages. The scan time was approximately 30 minutes. The distribution of migratory cells was manually labeled using MNI-Display software. (http://www.bic.mni.mcgill.ca/ServicesSoftwareVisualization/Display). The cells on the MRI images appear distinctly darker than white matter intensity and look similar to the intensity of gray matter. Previous studies had recognized a periventricular signal that was distinct from that of the overlying developing white matter (*42*).

*3D reconstruction of the brain*: To visualize the folded cortical surface of the term human (public open source from developing human brain project 35.7 GW at birth and 36.7 GW at scan (*43*)) and P0-aged piglet brain, we reconstructed the brains from the MRI images as a three-dimension using the ITK-SNAP software (*44*) and NEOCIVET pipeline (*45*). And we utilized the public open source for 4.5 years old Marmoset MRI (*46*) and P0-aged Mouse MRI (USCLA Laboratory of Neuroimaging). Briefly, the cortical surfaces were manually segmented, and segmented stacks were reconstructed for 3D visualization. To reconstruct the structures of the piglet Arc and lateral ventricle as a 3D, we combined the MRI images of P0-aged piglet brains and their serial Nissl-stained images. The Arc was defined as the cell-dense area near the ventricular wall, containing tiers 1, 2, and 3. To further match anatomies from MRI and Nissl-stained images, the entire structures, like the folding cortical morphologies, and the local landmarks including the striatum and hippocampus, were considered.

*Structural features analysis*

*Arc area measurement*: From the serial coronal Nissl-stained images of the human, piglet, macaque (public open source from NIH Blueprint NHP Atlas), marmoset, and mouse, we measured the Arc area and total brain area. To match the same anatomical location between species, we selected the serial coronal sections, ranging from the axis with the anterior striatum to the caudate nucleus, a part of the posterior striatum. To obtain the Arc area ratio (%), the Arc area was divided by the total brain area on the same slide. Tier 1 was measured as a cell-dense layer near the ventricular wall and tier2-3 area was the total Arc area minus the tier1 area.

*Gyrification Index*: The gyrification index (GI) from the animals was calculated using a series of coronal Nissl-stained images. Briefly, a contour connecting all cortical surfaces was drawn, and measured the length of the line. And the length of the entire contour connecting each sulcus and gyri was then divided by the length of the line connecting the cortical surface. To minimize the variation of the GIs across the species, we compared the GIs from the same range of the serial coronal sections containing from the anterior striatum to the caudate nucleus, a part of the posterior striatum. The GI from the mouse brain should be around 1.

*Quantification of a-SMA area*: To estimate the vascular areas within the Arc, alpha-smooth muscle actin (a-SMA) as a marker for vascular smooth muscle cells was used to label the vascular regions. Fluorescent images of a-SMA were processed to get the real signals by applying the threshold in the Fiji software. The apparent false-positive signals were manually filtered out by eye. The a-SMA-positive areas were calculated and divided by the area of tier1 and tier2-3 within the Arc.

Tiers thickness measurement: The boundaries between tier 1 and tiers2-3 were drawn based on the differential expression of DAPI and DCX protein. To measure the thickness of tiers from the ventricular wall, we measured the length in a radial straight line from the inside of the ventricular wall. The serial three slides from the different animals (n=3) were taken for quantification.

*Arc 3D volumetric measurement*: To estimate the volume of the piglet Arc across ages, we utilized the serial coronal sections of Nissl staining with 750 μm intervals. To match the anatomy of Arc from each age, each slide from different ages is matched based on the anatomical landmarks, such as the anterior striatum and caudate nucleus. Arc areas were measured in each section and followed equation 1. The total number of serial sections is *N*. For example, the Arc area in the first section is $A_{n=1}$, and the Arc area in the last section is $A_n$. The distance between the slides (h=750 μm).

$$\textbf{(1)}\ \text{Estimated volume} = \sum_{n=2}^{N}\left\{\left(A_{n-1} + A_n + \sqrt{A_{n-1}\cdot A_n}\right)\cdot\frac{h}{3}\right\}$$

*Histochemical staining*

*Cresyl staining (Nissl)*: All steps were performed at room temperature, unless otherwise specified. Sections were stained at a regular interval of 750μm. Frozen slides were allowed to thaw and equilibrate at room temperature overnight. Slides were baked for 20 mins at 60°C and incubated in cresyl violet solution (Sigma-Aldrich, cat# V5265) for 30 mins. Slides were washed with

distilled water twice before dehydration in an ethanol gradient. Slides were sequentially incubated in 50%, 70%, 95%, 100% ethanol in distilled water for 1 min each, followed by incubation in xylene solution for 3 mins. Slides were mounted and cured overnight and imaged using a Leica Aperio Versa 200 slide scanner microscope (Octopus) or Leica Widefield microscope.
*Immunohistochemistry*: All steps were performed at room temperature, unless otherwise specified. Frozen slides were thawed overnight at 4°C and allowed to equilibrate at room temperature for 3 hours. For antigen retrieval, slides were boiled at 95-100°C in 10 mM sodium citrate buffer (pH 6.0) for 5-10 mins, and then cooled at room temperature. Samples were permeabilized with 0.05 % Triton X-100 in PBS for 10 mins, then incubated in 1% $H_2O_2$ in PBS for 1 hour and blocked with TNB blocking buffer (0.1 M Tris-HCl; pH 7.5; 0.15 M NaCl; 0.5% blocking reagent from PerkinElmer, cat# FP1012) for 1 hour. Slides were incubated with primary antibodies in the TNB overnight at 4°C (see Table S4), followed by incubation with biotinylated secondary antibodies, diluted 1:250 in TNB solution for 2.5 hours at room temperature. Next, slides were incubated with horseradish peroxidase (HRP)-conjugated streptavidin, diluted 1:200 in TNB for 30 mins, before incubation with tyramide-conjugated fluorophores (Akoya), diluted 1:100, in amplification buffer (PerkinElmer) for 5 mins. Each dilution is the following, Cy3, Cy5, and FITC are 1:100, and Opal fluorophores are 1:50.

*RNAscope and protein co-detection*: RNAscope probes and reagents were obtained from Advanced Cell Diagnostics (ACD). In preparation for mRNA detection, slides were removed from -80°C, baked at 60°C for 30 mins, and then postfixed in 4% PFA for 15 mins at room temperature. Next, slides were sequentially dehydrated in 50%, 70%, and 100% ethanol in distilled water for 5 mins each, and fully dried at room temperature. This was followed by incubation in 3% $H_2O_2$ in PBS for 10 mins and a wash with distilled water, before antigen retrieval (ACD, Co-detection Target Retrieval Reagent, cat# 323180) for 5 mins at 98-102°C. Slides were quickly washed in distilled water and PBS-T (0.1% Tween 20 in PBS), and incubated with DCX primary antibody, diluted 1:200 in ACD diluent (Co-detection Antibody Diluent). Slides were incubated with primary antibodies diluted in ACD diluent (Co-detection Antibody Diluent), overnight at 4°C. Slides were washed with PBS-T and fixed with 4% PFA for 30 mins at room temperature, which was followed by incubation with protease III for 20 mins at 40°C in the RNA scope oven (HybEZ, ACD) and a wash in distilled water. Probe hybridization and amplification steps were performed using the RNAscope HiPlex v2 kit (ACD, cat# 324419). Slides were incubated with the desired probes for 2 hours at 40°C in the hybridization oven and washed with a wash buffer. For amplification steps, slides were incubated with Amp1 for 30 mins at 40°C and washed twice in the wash buffer for 2 mins each. Next, slides were incubated with Amp2 for 30 mins at 40°C, and then washed twice in the wash buffer, followed by Amp3 buffer for 30 mins at 40°C and two washes. For fluorophore labeling, slides were immersed in RNAscope Hiplex Fluoro buffer and incubated for 15 mins at 40°C. Slides were washed with a wash buffer, and incubated in biotinylated secondary antibodies, diluted 1:250 in ACD diluent, for 30 mins at room temperature. Slides were washed with PBS-T for 2 mins twice and incubated with HRP-conjugated streptavidin, 1:200 in ACD diluent, for 30 mins at room temperature. Finally, slides were incubated with tyramide-conjugated Opal 690, 1:50, for 10 mins at room temperature. Slide was added of DAPI solution for 30 sec at RT and mounted with ProLong Gold Antifade Mountant solution (Thermo Fisher Scientific, cat# P36934).

*HiPlex FISH*: For the multiplex fluorescent Hiplex Assay (v2, ACD), slides were prepared in the same way as for RNAscope and protein co-detection experiments. The HiPlex assay allows multiplexed detection of up to 32 targets on the same tissue sample (see Table S5). Briefly, sections were baked at 60°C, fixed in 4% PFA, dehydrated with 50%, 70%, and 100% ethanol, exposed to antigen retrieval, and incubated with protease III for 20 mins at 40°C. For the first hybridization cycle (cycle 1), 12 target probes were hybridized, and amplified together. Since the fluorophores are three (T1-T3: 488, 555, 647), which can be detected together, 1 cycle (12 probes) consists of 4 rounds of fluorescence detection. Slides were imaged as specified below. After imaging, coverslips were gently removed after soaking slides in 4X SSC buffer (diluted in distilled water from 20X SSC stock; Invitrogen, cat# 15557044) for at least 1 hour at room temperature. Slides were treated with 10% cleaving solution v2 (ACD) for 15 mins at room temperature to cleave off the conjugated fluorophores from the previous round. The second round of conjugation of fluorophores was repeated in the same way. Before the second hybridization of the cycle (cycle 2), slides were incubated with HiPlex Up Reagent (ACD) for 5 mins at room temperature and washed with PBS-T. This step was repeated three times. The hybridization of probes, amplification, and detection were the same as those of the first cycle. A third hybridization cycle (cycle 3) with a final set of 9 target probes was completed as before. Images were acquired using a Leica Stellaris 8 Tau STED confocal Microscope with a 40X objective (HC PL APO 40X/1.30 Oil CS2). To obtain the exact locations in each round, several landmarks were labeled on each slide. Images from all 13 rounds were registered using HiPlex image registration software v2 (ACD).

*Organotypic slice cultures and live imaging*: The fresh piglet brains at P0-aged were cut in a coronal block covering the cingulate and piriform cortex in cold artificial cerebrospinal fluid (ACSF), which had been oxygenated for at least 1 hour. ACSF contained 125 mM NaCl, 2.5 mM KCl, 1 mM $MgCl_2$, 1 mM $CaCl_2$, and 1.25 mM $NaH_2PO_4$. Brains were embedded in 3.5% low-melting-point agarose (Fisher, cat# BP165-25) and sectioned using a Leica VT1200S vibrating blade microtome as 300µm slices with the following operational settings: speed, 2mm/s; amplification 0.8mm. Slices were transferred into Millicell-CM slice culture inserts (Millipore, cat# PICM03050) that were immersed in the modified DMEM/F12+Glutamax™-I culture medium (Gibco, cat#10565018) with 1X N2 Supplement (Gibco, cat#17502048), 0.05X B27 Supplement (Thermo Fisher, cat#17504044), 20ng/ml hFGF-2 (Gibco, cat#13256029), 20ng/ml hEGF (Gibco, cat#PHG0311), 20µg/ml human insulin (Sigma, cat#I9278), 5ng/ml BDNF (Sigma, cat#SRP3014), 10µM Rock inhibitor Y-27632 (Stemgent, cat#04-0012), 100U/ml Penicillin/Streptomycin (Gibco, cat#15140-122). Tissue slices were microinjected with adenovirus (AV-CMV-GFP, $1x10^{10}$ PFU/ml, 1-2 µl; Vector Biolabs) into areas of interest and were kept at 37°C with % $CO_2$ and, 8% $O_2$. For time-lapse imaging, the media was changed into the modified Basal Medium Eagle (BME; Gibco, cat#21010046) with 25% Hanks' Balanced Salt Solution (HBSS; Gibco, cat#14025092), 5% fetal bovine serum (FBS; Gibco, cat#10437028), 1% N2 supplement, 0.66% D-(+)-glucose (Sigma, cat#G7528), and 1% Penicillin/Streptomycin. Samples were loaded into a 37°C chamber of an inverted Leica Stellaris 8 Tau STED microscope with an on-stage incubator streaming 5% $CO_2$. Samples were imaged using a 10X air objective at 25 mins intervals for 72 hours with repositioning of the z-stacks every 8-9 hours. For post hoc immunostaining, samples were fixed in 4% PFA and immunohistochemistry was performed as described above. For long-term tracing of GFP+ cells, samples were cultured for 21 DIV and sequentially fixed at 12 DIV and 21 DIV, respectively. Half of the cultural media was replaced with fresh media every 2 days.

*Tissue clarification:* Fixed piglet brains at P0-5 age were cut coronally into 3mm thick tissue blocks. Reagents for tissue clearing were obtained from Binaree Inc. Blocks were sequentially incubated with tissue Clearing Solution A and B in a rotator for 3 days each at 37°C. Blocks were washed with distilled water with constant agitation at 4°C for 2 hours, and permeabilized with a permeabilization buffer (0.3% Triton X-100, 10% DMSO, and 5% BSA (bovine serum albumin) in PBS) for 3 days at 37°C. The primary antibody (DCX, 1:500) was diluted in a blocking buffer (0.5% Tween 20, 5% DMSO, and 5% BSA in PBS) and incubated for 3 days at 37°C. Blocks were washed with PBS-T (0.1% Tween 20) under constant agitation at 4°C for 3 hours, followed by incubation with the secondary antibody (Alexa 488, 1:500) for 3 days at 37°C. Blocks were washed with PBS-T and incubated within Mounting & Storage Solution (Refractive Index, 1.46) in a shaking incubator at 37°C for 1 day. Images were acquired using a Light-sheet Microscope (AZ-100) and 2-5X magnification.

*Imaging, processing, and quantification*

*Histological imaging analysis:* Tiled images of entire slides were acquired on a Ziess Widefield or Keyence BZ-X Analyzer at objective 10X (NA 0.45) and files were stitched automatically. Higher magnification of images was acquired on a Stellaris confocal microscope using 10X (NA 0.3) 40X, and 63X objectives. Images used for quantification were acquired with a total of 3 replicates (slides) for each animal (5 cases of human samples, 3 cases of piglet at each age, 3 cases of marmoset samples, and 2 cases of mouse samples). Images were processed in Fiji where adjustments to image brightness and contrast were made. Further, quantifications were performed in Neurolucida software (MBF Bioscience, 2018 version) and cells were manually counted throughout the z-stack images for all quantifications. Quantifications were performed and repeated by three authors.

*Quantification of DCX expression intensity within the Arc:* For the measurement of DCX immunostaining signal intensity from tiers 1 to 3, fluorescent images of DCX+ were processed to obtain real signals by applying the threshold in the Fiji software. The apparent false-positive signals were manually filtered out by eye. The signal intensity of DCX-positive pixels was calculated in a rectangular box containing from the ventricular wall to tier 3 regions. The boundary between tiers was determined based on the DAPI signals. The serial three slides from the different animals (n=3) were taken for quantification.

*Quantification of DCX+ cells expressing the transcription factors*: Images used for quantifications were acquired covering the entire tiered Arc region. Based on the differential expression of DAPI and DCX protein, tiers 1, 2, and 3 were isolated. 4-5 images per tier (total 12-15 images per slide) were acquired with a thickness of 0.22 micron for each z-stack, with a total of 3 replicates (slides) for each animal. Images were processed on ImageJ where adjustments to image brightness and contrasts were made. Cells were manually counted throughout the z-stack images for all quantification. Quantifications were performed and repeated by three authors. For a heatmap of the enrichment of subtypes of DCX+ cells across tiers, the average values of subtype's proportion in each tier were utilized. R studio was used to compare the relative expression patterns of each subtype across tiers.

*Mapping of DCX+ cells across species*: Human, piglet, and marmoset brains were sectioned at regular intervals of 750µm along the anterior and posterior (A-P) axis. To match the A-P position of sections across species, anatomical landmarks were utilized, such as the striatum, caudate nucleus, thalamus, and temporal gyrus, Slides were stained with anti-DCX primary antibody (1:200) to visualize DCX+ streams across brains and imaged at 10X on a Zeiss Axiovert 200M Microscope. Neurolucida software (MBF bioscience) was used for image collection and analysis. For each DCX+ cell, the soma with the leading neuronal process was marked.

*HiPlex FISH image processing*: To quantify transcript abundance from HiPlex FISH images, regions of interest were drawn around *DCX*-positive nuclei. Next, images were processed with the Reyni Entropy filter, resulting in a binary mask. The expression value of the transcript was then computed as the pixel percent area of the probe signal in the mask in reference to the ROI. This process was repeated across all genes in the probe set, and a cell by gene expression matrix was generated. After visual analysis, we determined that only cells with greater than the median expression level for a particular gene demonstrated true expression of that gene. These gene expression values below the median were treated as background and set to zero for that cell. For *COUP-TFII* and *SP8*, the cutoff was set at a percentile threshold of 40% based on their more robust expression. The resulting expression matrix was loaded as a Seurat R object. Expression values were log-normalized and scaled before conducting principal component analysis. All principal components were used to conduct Leiden Neighborhood clustering and produce UMAP embeddings.

*Neighborhood analysis*: The interaction score from the histoCAT neighborhood analysis (*47*) was adapted to quantify statistically significant interactions that occurred between cell types. Using the cell-type classifications from the spatial transcriptomic analysis, interactions were quantified by comparing the distances between all cells in each image. Cells with distances less than 4 pixels were defined as neighbors. The number of pairwise interactions between cells of the same type and different types was quantified and then compared to a distribution generated from the same image with randomized assignments of labels. We then conducted a one-tailed permutation test to produce a p-value, which represents the likelihood of neighborhood interaction compared to the randomized distribution. To visualize the interactions between cell types, we use the present cell interaction graph. We only visualize interactions if their interaction is significant ($P < 0.05$) in at least 30% of the images and simultaneously present in at least 90% of the images.

*Statistical Analysis*: Standard errors were shown for cell quantification. Pearson's Correlation analysis and MANOVA analysis were performed in GraphPad Prism (v.6).

**Supplementary Text**

**Figure S1.**

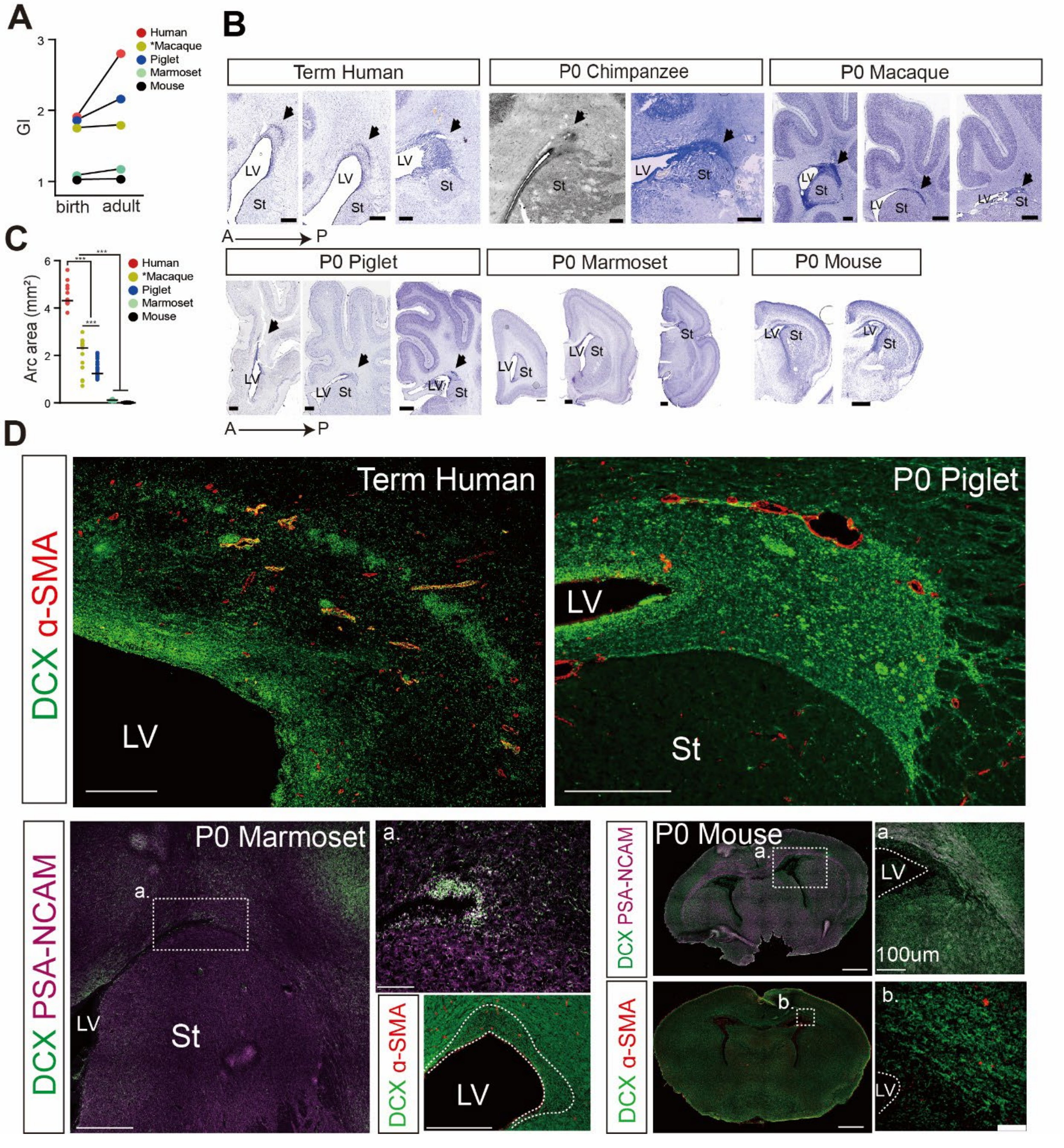


**Fig. S1. Comparative analysis of ventricular wall structure across species.**

**A.** Developmental changes in cortical folding as measured by Gyrification Index (GI) across species. Human, macaque, and pig brains exhibit gyrencephalic features, while marmoset and mouse brains possess smooth surfaces, even in adults. The Macaque dataset is from the public open source (marked with *). Humans and marmosets are both primates and shared a last common ancestor approximately 40 million years ago. Mice are rodents, which shared a last common ancestor in the Euarchontoglires clade with primates approximately 85 million years ago. Pigs are in the more distant Laurasiatheria clade (with bats, horses, dogs, cats, and whales) and shared a last common ancestor with euarchontoglires approximately 95 million years ago (*14*).

**B.** Nissl-stained Serial coronal brain sections from different species collected at birth. The enlarged SVZ, termed the Arc, is present in human, chimpanzee, macaque, and piglet brains, but not in marmoset and mouse brains. Scale bar, 500µm (human, chimpanzee, macaque, piglet, marmoset); 100µm (mouse). Lateral ventricle (LV); striatum (St).

**C**. Quantification of Arc area ($mm^2$) from Nissl-stained sections of different species. The Macaque dataset is from the public open source (marked with *).

**D.** Confocal images show robust expression of the migratory neuron marker DCX (in green), and an abundance of blood vessels expressing α-SMA (in red) in the neonatal human and piglet Arc. Migratory neural populations expressing DCX and PSA-NCAM (in magenta) do not form a tiered structure and vascular areas are sparse in the ventricular wall of P0 marmoset and mouse brains. Scale bar, 500µm; 100um (higher magnification images). Lateral ventricle (LV); striatum (St).

#### Figure S2.

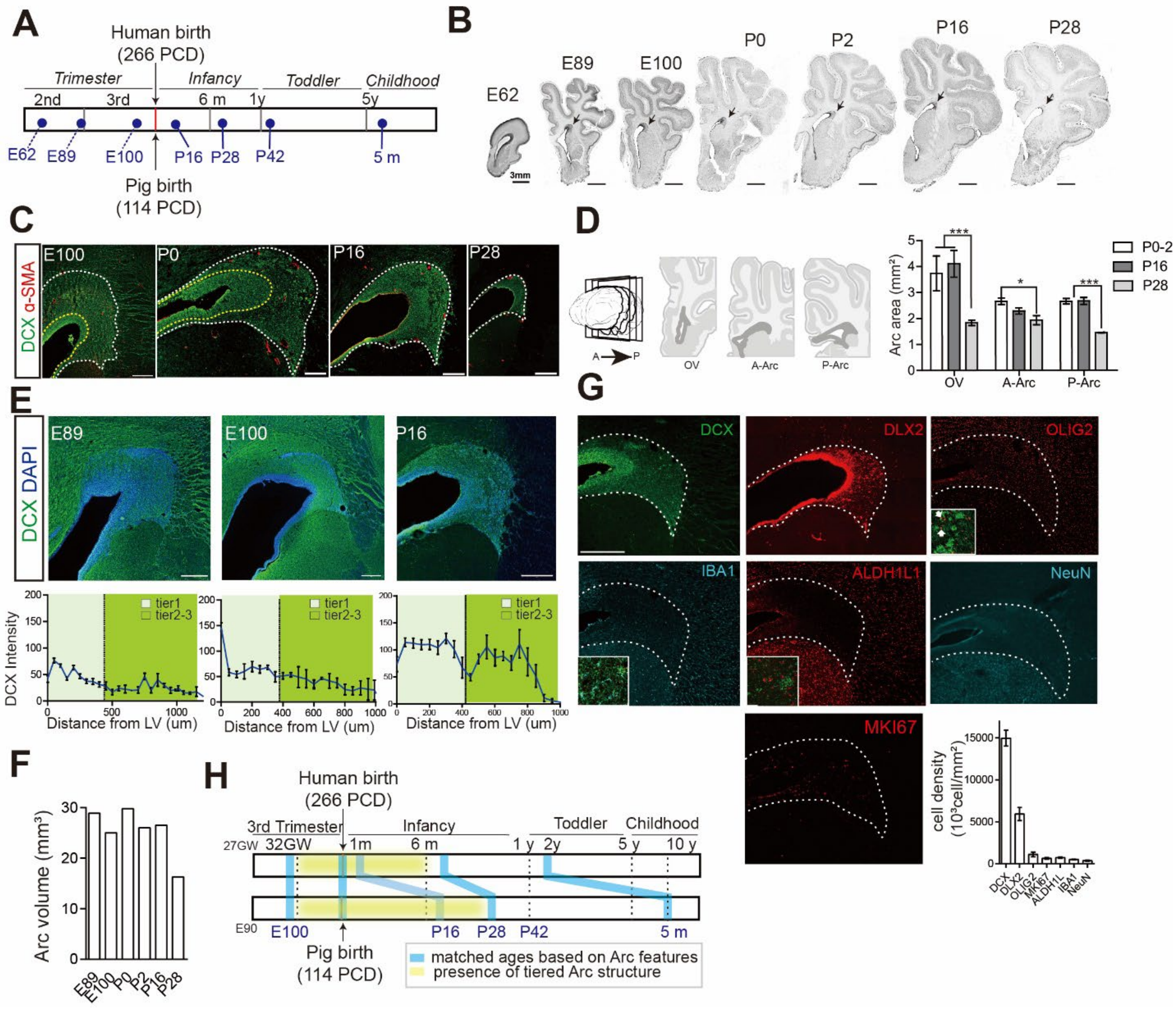


**Fig. S2. Structural changes in the piglet Arc during perinatal stages.**

**A.** Schematic showing the stages of human (top) and matched pig (bottom) development (*10, 11, 48*). Dashed lines indicate predictions of corresponding stages along the pig developmental timeline when compared to human anatomical development. PCD, post-conception day.

**B.** Nissl-stain of coronal section from piglet brains across pre- and postnatal ages. The Arc (arrow) is clearly present up until P16. Scale bar, 3mm.

**C.** Confocal images of core Arc structural changes during perinatal stages of the pig. Tier1 is characterized by a prominent population of DCX+ neurons. α-SMA+ vasculature at E100 and P0. are clearly observed. These features disappear by P16 and P28. The yellow dashed line highlights the boundary between tiers 1 and 2; the white dashed line between tiers 3 and 4. Scale bar, 500µm.

**D.** Left: Schematic showing piglet Arc subregions at three selected coronal planes along the anterior (A) to posterior (P) axis, illustrated for P0 planes cover the olfactory ventricle (OV), anterior Arc (A-Arc), and posterior Arc (P-Arc). Right: Quantification of Arc area at each plane across ages. The Arc area along the A-P axis is significantly reduced at P28. Data means ± SEM (

*, $p < 0.05$; **, $p < 0.01$; ***, $p < 0.001$ by unpaired t test).

**E.** Quantification of DCX expression intensity across tiers in E89, E100, and P16 piglet brains. Tier 1 is separated from tiers 2-3 based on DAPI intensity (black dashed line). The tiered structures, imposed by differential expression of DCX, are not yet established at embryonic stages. Lateral ventricle (LV). Scale bar, 500µm. Data means ± SEM.

**F.** The 3D volume of the Arc was measured from Nissl-stained piglet serial sections (coronal, with 750 um intervals). To match the anatomy of the Arc across ages, sections were matched based on anatomical landmarks, such as the striatum and caudate nucleus. The Arc volume decreased by up to 40%, from P16 to P28.

**G.** DCX+ and DLX2+ cells are abundant in the P0 piglet Arc (dashed white line), compared to other cell types, such as oligodendrocytes (marked by OLIG2), microglia (IBA1), astrocytes (ALDH1L1), and mature neuron (NeuN), as well as proliferative cells (MKI67). The density of each population is measured within the total Arc area. Scale bar, 500µm. Data means ± SEM.

**H.** Schematic showing Arc features age-matched between humans and piglets (blue lines), as well as the period when the tiered Arc structure is present in human and piglet brains (highlighted in yellow). Gestational week (GW).

### Figure S3.

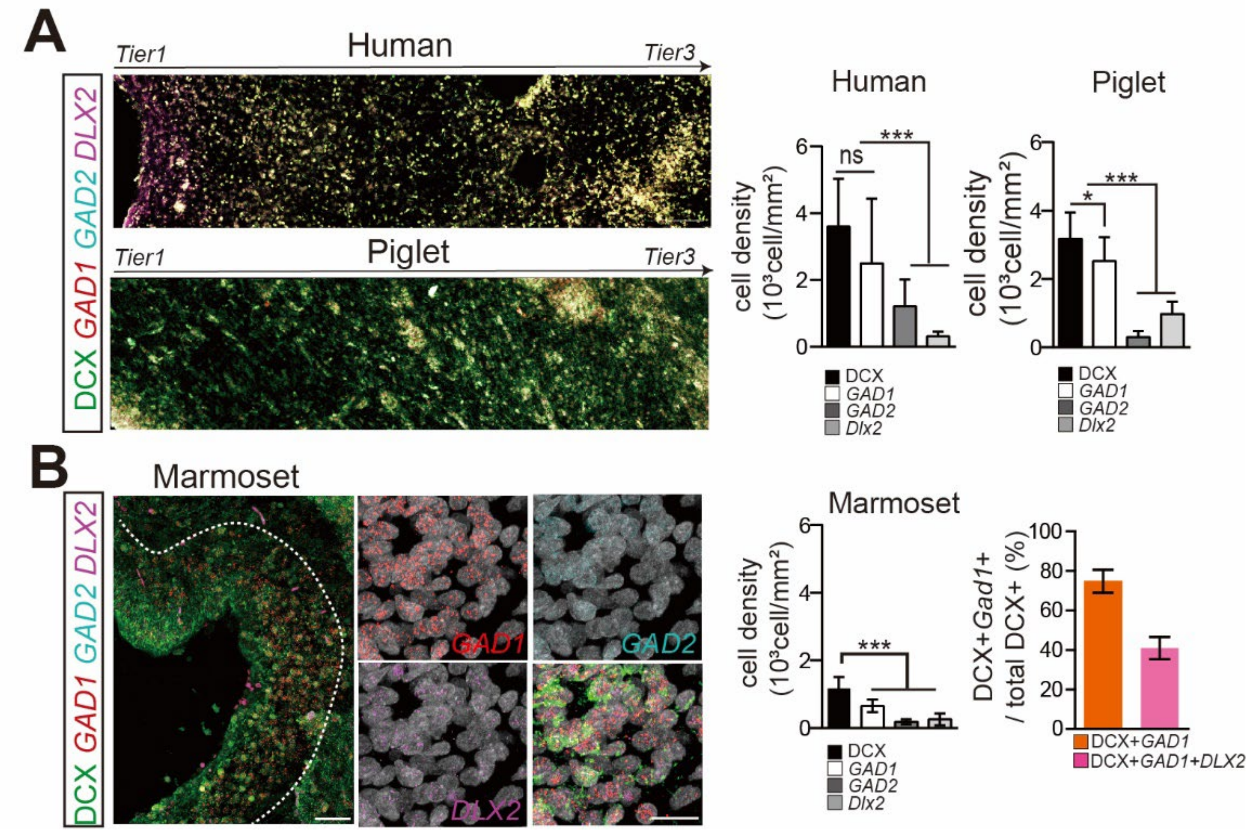


**Fig. S3. Cellular composition within the Arc across species.**

**A.** Left: Co-detection of mRNAs for GABAergic interneuron markers and DCX protein across the tiered Arc on the neonatal human and P0 piglet brains. Scale bar, 30µm. Right: Quantification of cell densities within the Arc. Data means ± SEM (*, $p < 0.05$; ***, $p < 0.001$ by unpaired t-test).
**B.** Left: Co-detection of mRNAs for GABAergic interneuron markers and DCX protein in the P0 marmoset ventricular wall. The white dotted line delineates the border of cell-dense regions of the SVZ. Scale bar, 100µm, and 15µm (higher magnification images). Right: Quantification of cell densities within the SVZ. Data means ± SEM (***, $p < 0.001$ by unpaired t-test).

#### Figure S4.

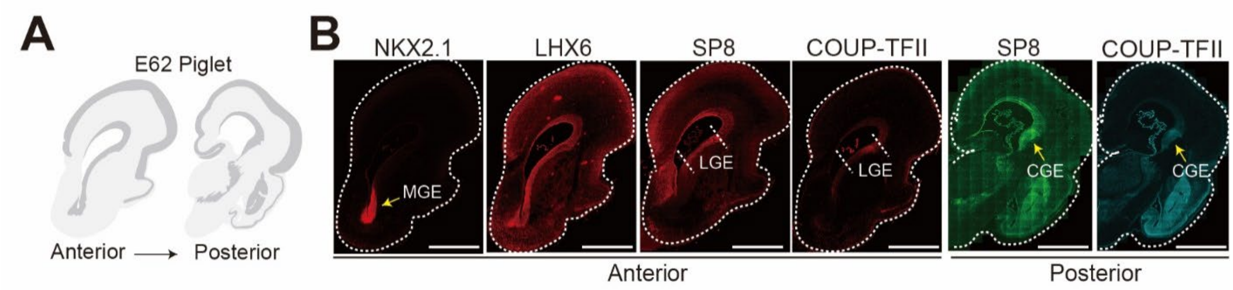


**Fig. S4. GE-associated transcription factors in the E62 piglet brain.**

**A.** Schematic illustrating anterior and posterior coronal sections of an E62 piglet brain, at planes shown in (B).

**B.** Wide-field images of an E62 piglet brain, at anterior (left) and posterior (right) planes. Coronal sections were immunostained with antibodies against selected transcription factors. Yellow arrows indicate areas of ganglionic eminence (GE) with increased expression of respective transcription factors. The medial GE (MGE) expresses exclusively NKX2.1, as shown for anterior sections. The caudal GE (CGE) has a stronger expression of SP8 and COUP-TFII, as shown from a posterior section. SP8- and LHX6-expressing cells are observed in the lateral GE (LGE), as shown for an anterior section; LHX6+ cells in the LGE are likely MGE-derived migratory neurons. Scale bar, 3mm.

**Figure S5.**

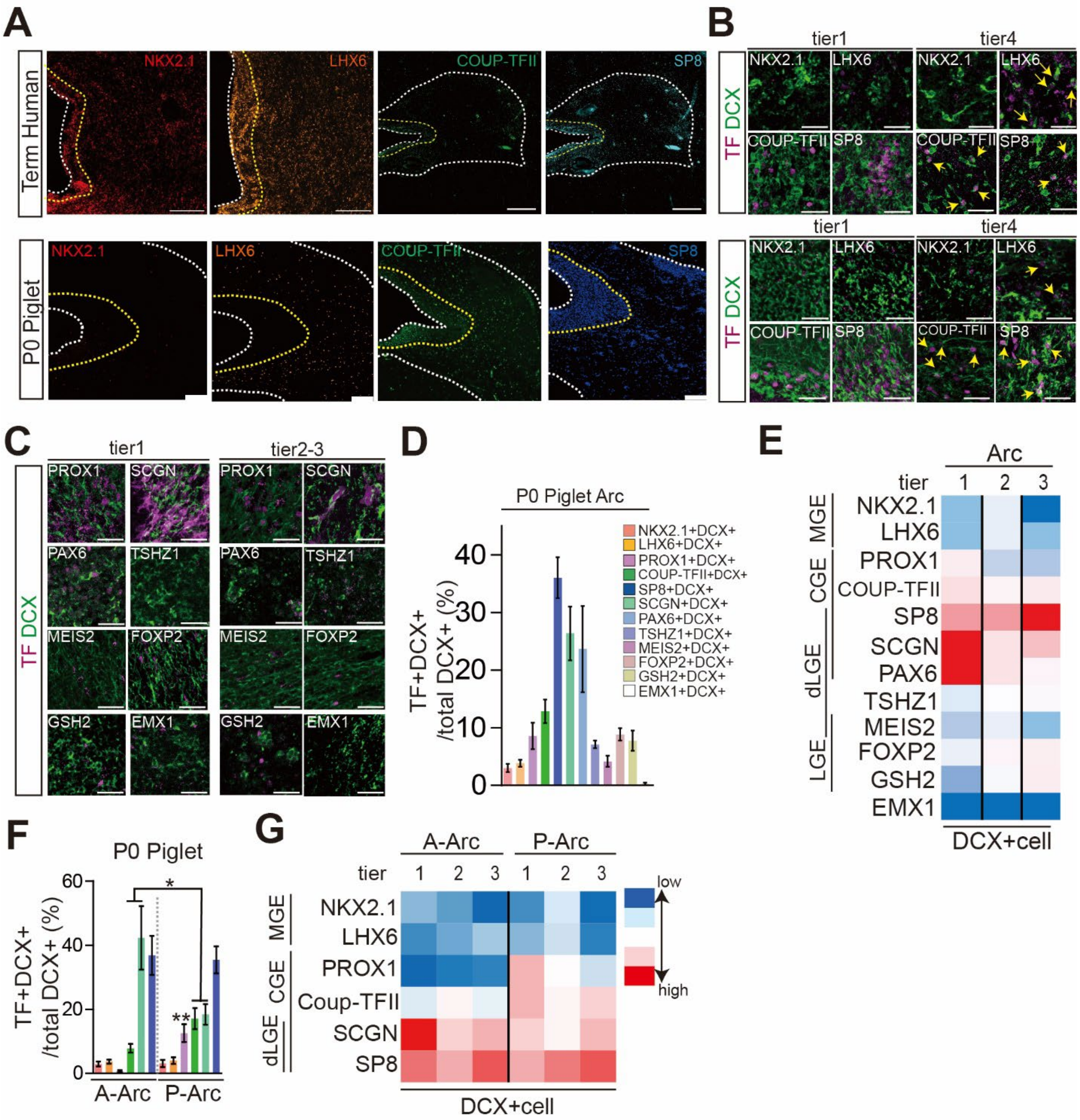


**Fig. S5. Diverse GABAergic interneurons within the Arc.**

**A.** Wide-field images of the neonatal human and P0 piglet Arc, immunostained with antibodies against GE-associated transcription factors. The yellow dashed line highlights the boundary between tiers 1 and 2; the white dashed line between tiers 3 and 4 (outer) or between tier 1 and the lateral ventricle (inner). Scale bar, 100μm.

**B.** Higher magnification images of the Arc show that DCX+ cells, which express the transcription factors (TF) SP8 and COUP-TFII, are abundant within the Arc, especially in tier 1 and 4, in both

term human (top) and P0 piglet (bottom). LHX6+DCX+ cells are more abundant in the human tier 4 than in the piglet tier 4. Yellow arrows indicate DCX+ cells expressing TFs. Scale bar, 20μm.
**C.** Higher magnification images of the P0 piglet Arc, immunostained with antibodies against additional TFs enriched in the GE. PROX1, SCGN, and PAX6 are associated with the caudal GE (CGE) and dorsolateral GE (dLGE) and are predominantly expressed by DCX+ cells in tier1. LGE-related TSHZ1, MEIS2, FOXP2, and GSH2 are rarely expressed by DCX+ cells within the Arc. EMX1 expression, associated with excitatory progenitor identity, is rare. Scale bar, 15μm.
**D.** Quantification of subpopulations of DCX+ cells in the P0 piglet Arc expressing different TFs enriched in GE. Data means ± SEM of counts performed on three individual cases (n=3).
**E.** Heatmap depicting the spatial distribution of DCX+ cells expressing different TFs across the tiers of the P0 piglet Arc. The color gradient represents TF expression levels from low (blue) to high (red). Medial GE (MGE).
**F and G.** Quantification of the subpopulations of DCX+ cells across anterior-posterior divisions of the Arc in the P0 piglet brain (F), and corresponding heatmap (G). PROX1+ and COUP-TFII+DCX+ cells, associated with CGE, are most abundant in the posterior Arc (P-Arc), especially in tier 1. SCGN+DCX+ cells are most abundant in tier1 of the anterior Arc (A-Arc). SP8+DCX+ cells are more evenly distributed across the entire Arc.

#### Figure S6.

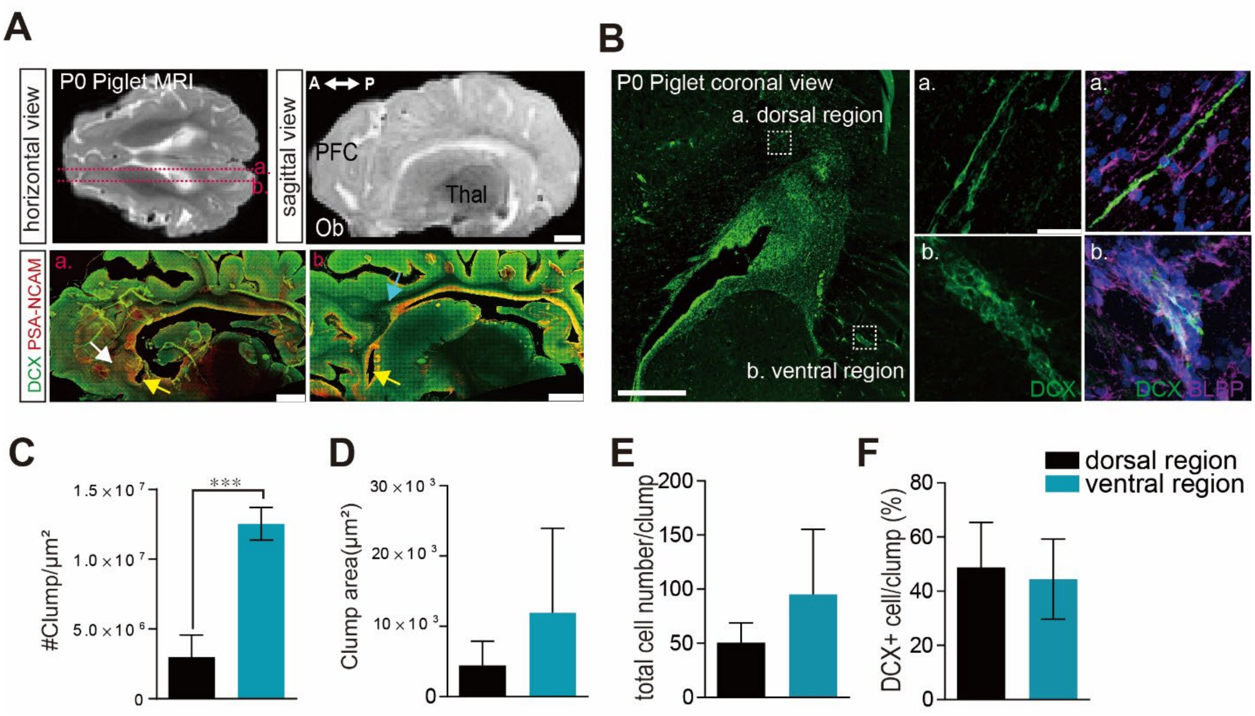


**Fig. S6. Multiple migratory streams emerge from the Arc.**

**A.** Representative T2-weighted MRI images of a P0 Piglet brain (top, left horizontal view; top, right: sagittal view). Bottom panels show the corresponding sagittal sections immunostained with antibodies against DCX and PSA-NCAM. The medial section (a) shows extensive bands from the opened olfactory ventricle (OV, yellow arrow). The lateral section (b) shows the enlarged SVZ (Arc) extending into the prefrontal cortex (PFC; blue arrow) as well as OV (yellow arrow). Scale bar, 500μm. Ob; olfactory bulb; Thal: thalamus.

**B.** Confocal images of a P0 Piglet coronal section, immunostained with antibodies against DCX and BLBP. The dorsal region (a) shows DCX+ cells migrating individually out from the Arc, with little association with BLBP+ cells. The ventral region (b) shows cell clumps of DCX+ cells migrating out from the Arc, surrounded by BLBP+ cells. Scale bar, 500μm, and 30μm (higher magnification images).

**C.** Quantification of the clump density in the dorsal and ventral regions adjacent to the P0 piglet Arc. The aggregation of three or more cells is defined as a clump. Ventral regions are significantly more enriched for cell clumps compared to dorsal regions. Data means ± SEM (***, $p < 0.001$ by unpaired t-test).

**D to F.** Profiling of clump features in the P0 piglet. Quantification of the clump area (D), the total number of DAPI+ nuclei in a clump (E), and the proportion of DCX+ cells in a clump (F). Clump area and size vary in the dorsal and ventral regions, but the proportion of DCX+ cells is similar in both regions at around 50%.

#### Figure S7.

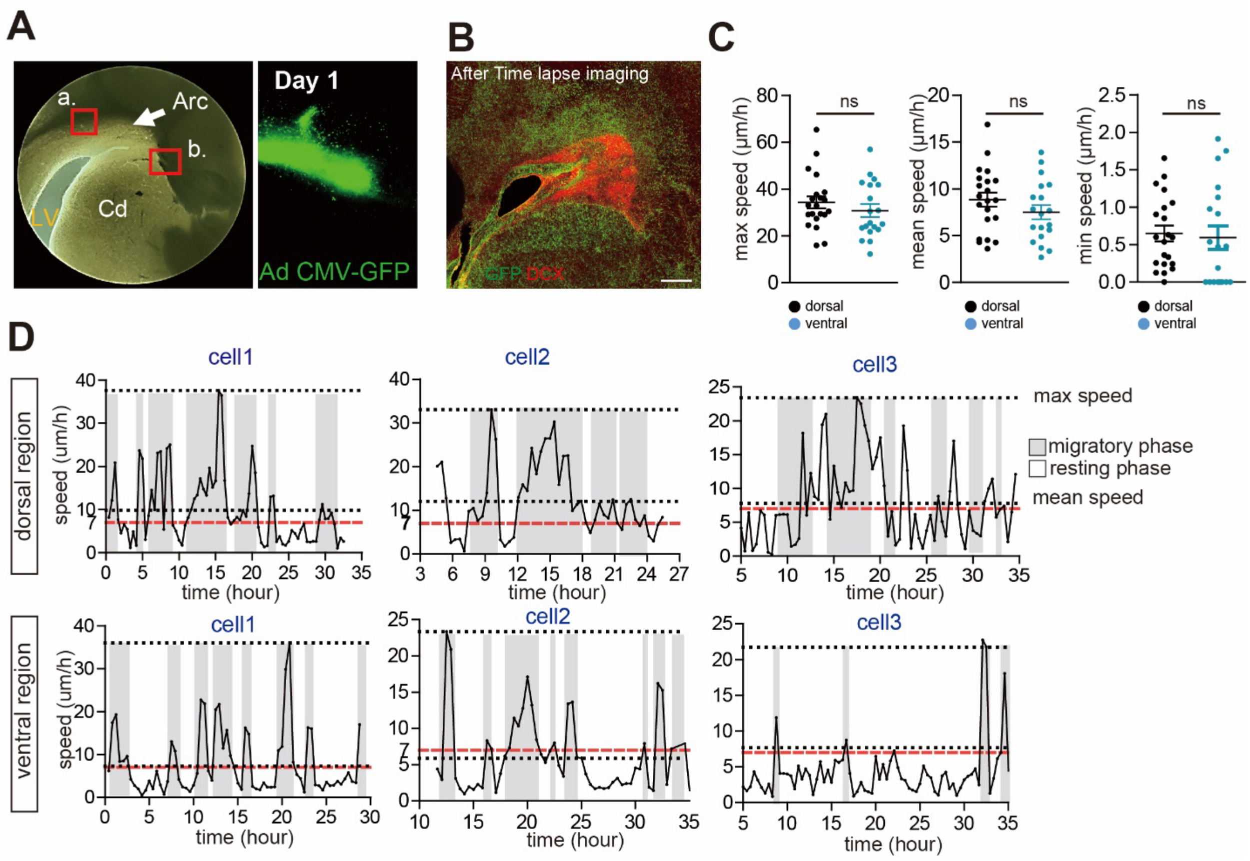


**Fig. S7. Time-lapse imaging of P0 piglet organotypic slice cultures.**

**A.** Left: bright-field images of a P0 piglet organotypic slice, containing the Arc (arrow). Boxed areas indicate dorsal (a), and ventral (b) regions analyzed in C and D. Right: GFP+ cells are present in the Arc on day 1 after microinjection of Adenovirus (Ad) CMV-GFP into the Arc. Lateral ventricle (LV); caudate nucleus (Cd).

**B.** Post hoc immunostaining after time-lapse imaging reveals that GFP+ cells within the Arc are mostly DCX+ and migrate out from the Arc into the dorsal and ventral areas. Scale bar, 100μm.

**C.** Quantification of the migratory speed of GFP+ cells from the dorsal and ventral regions. No significant difference in maximum, mean, and minimum speeds was observed. Data means ± SEM.

**D.** Speed profile of cells over time, with a maximum speed (black dashed line, top) that is much higher than the mean speed (black dashed line, bottom). Migratory phases (soma speed higher than 7μm/h; red dashed line) are indicated as a gray box and resting phases (soma speed less than 7μm/h) as a white box. Three representative cells each from the dorsal and ventral regions are shown. Migrating cells in the ventral regions exhibit more heterogeneity in their migratory behaviors than those in the dorsal regions.

**Figure S8.**

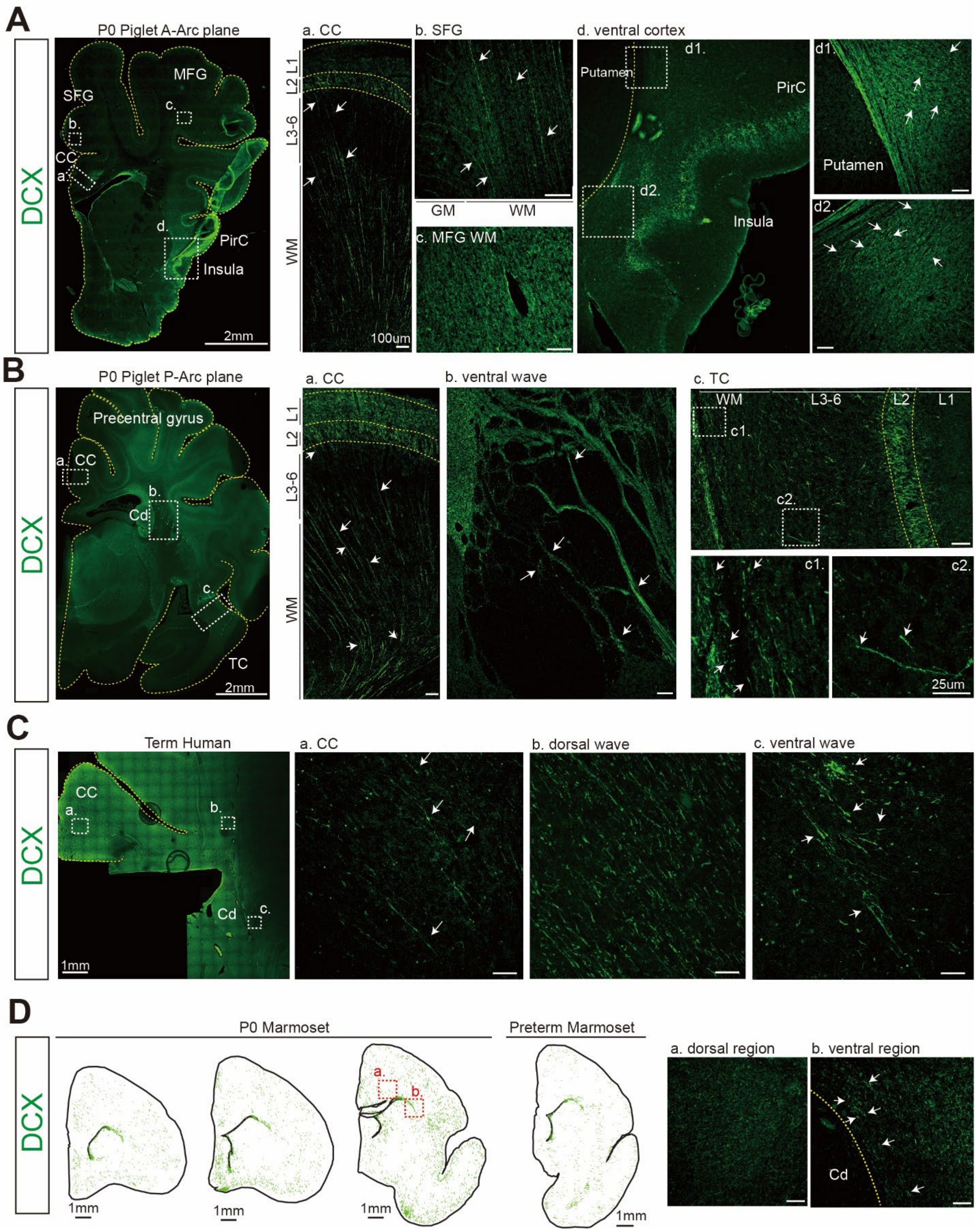
A
DCX
P0 Piglet A-Arc plane
MFG
SFG
CC
PirC
Insula
2mm
a. CC
L1
L2
L3-6
WM
100um
b. SFG
GM
WM
c. MFG WM
d. ventral cortex
Putamen
PirC
Insula
d1.
d2.
B
P0 Piglet P-Arc plane
Precentral gyrus
Cd
TC
2mm
b. ventral wave
c. TC
c1.
c2.
25um
C
Term Human
1mm
b. dorsal wave
c. ventral wave
D
P0 Marmoset
Preterm Marmoset
1mm
a. dorsal region
b. ventral region

**Fig. S8. Dorsal and ventral migratory streams from the Arc target distinct cortical regions, associated with higher cognitive function, in gyrencephalic brains.**

**A.** Coronal section of the P0 piglet brain, at the level of the anterior Arc (A-Arc; refer to fig. S2D), immunostained for DCX. (a to d) Higher magnifications of the boxed areas in (A, left) show that DCX+ cells migrate out from the Arc to the cingulate cortex (CC, a), superior frontal gyrus (SFG, b), and ventral cortex (piriform cortex (PirC) and insula; d) but less so to the middle frontal gyrus (MFG, c). Arrows indicate DCX+ cells with elongated, migratory morphology. Scale bar, 2mm (A, left), 100µm (a and d), and 50µm (b, c, d1, d2). Gray matter (GM); layer (L); white matter (WM).

**B.** Coronal section of the P0 piglet brain, at the level of the posterior Arc (P-Arc; refer to fig. S2D), immunostained for DCX. (a to c) Higher magnifications of the boxed areas in (B, left). DCX+ cells migrate into the posterior CC (a) and temporal cortex (TC, c). Arrows indicate DCX+ cells with elongated, migratory morphology. Scale bar, 2mm (B, left), 100µm (a to c), and 25µm (c1 and c2). Caudate nucleus (Cd).

**C.** Coronal section of a term human brain, immunostained for DCX at a plane equivalent to the P-Arc plane of the P0 piglet brain illustrated in (B). (a and b) DCX+ cells appear individually with an elongated morphology (arrows), migrate into the posterior CC (a), and are densely packed in the dorsal white matter (b). (c) DCX+ cells near the ventral Cd exist in clumps. Scale bar, 1mm (C, left), 50µm (a to c).

**D.** Spatial mapping of DCX+ cells (in green) in serial coronal sections of P0 and a preterm marmoset brain; few DCX+ cells (arrows) are present in the dorsal (a) and ventral side (b) of the ventricular wall. Scale bar, 1mm (map), 30µm (confocal images).

**Figure S9.**

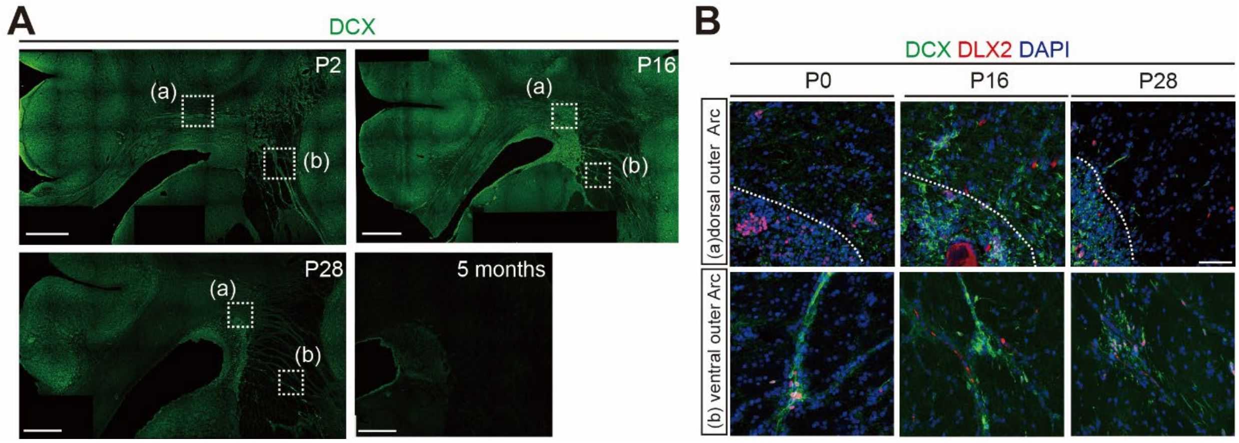


**Fig. S9. Developmental timing of dorsal and ventral migratory streams from the Arc.**

**A.** Wide-field images showing the distribution of DCX+ cells across postnatal stages of the pig. Dorsal streams are detectable until P16, and ventral streams are observed until P28. All streams disappear by 5 months after birth. Higher magnifications of the boxed areas in (A) were analyzed in (B). Dorsal outer Arc region (a) and ventral outer Arc region (b). Scale bar, 500µm.

**B.** Confocal images of DCX+DLX2+ cells in the dorsal and ventral outer Arc of P0, P16, and P28 piglet brains. Migration of DCX+ cells from the dorsal outer Arc peak at P16 and decreases thereafter. White dashed line indicates a boundary between Arc and dorsal outer Arc. Migration of DCX+ cells from the ventral outer Arc persists until P28. Scale bar, 50µm.

#### Figure S10.

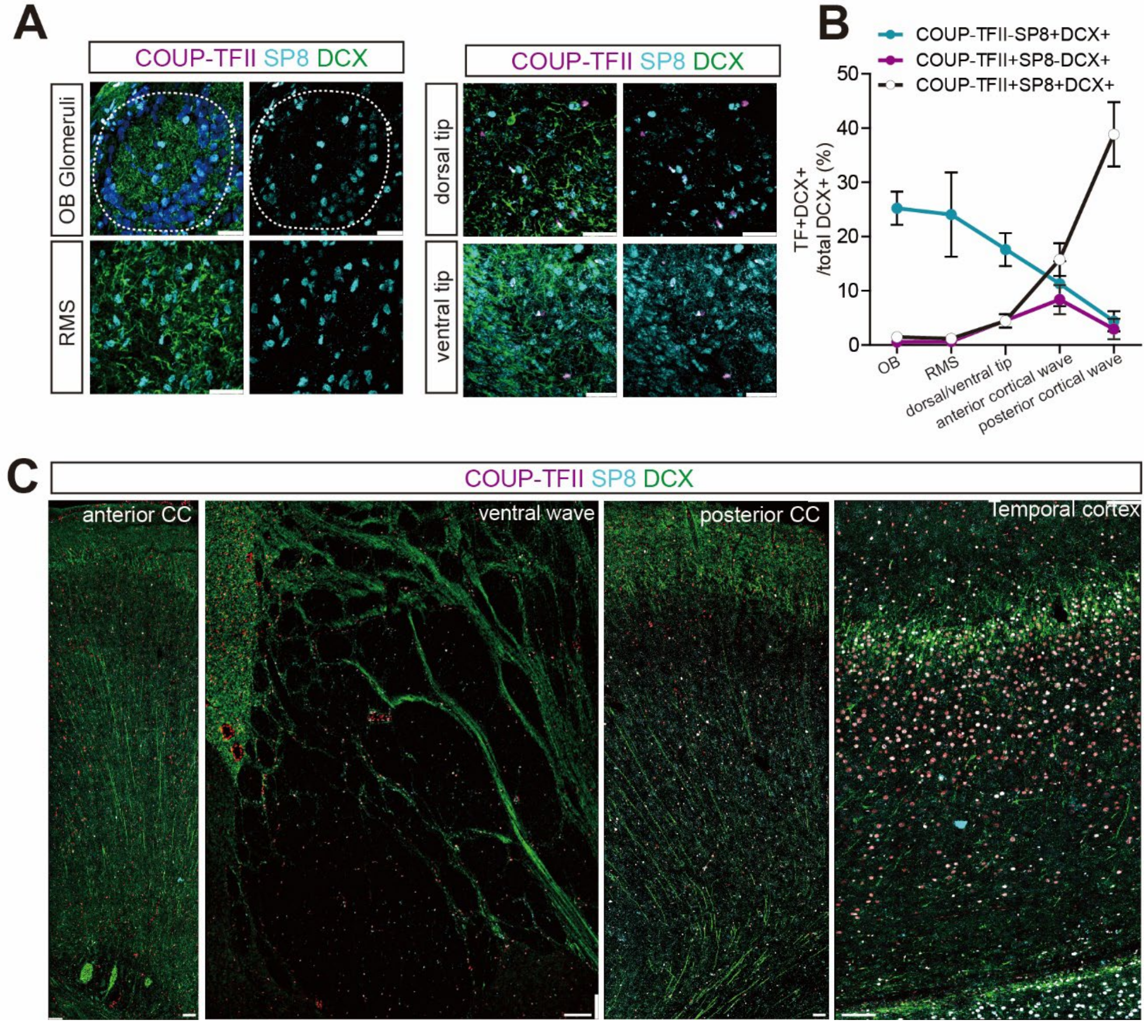


**Fig. S10. CGE-derived COUP-TFII+SP8+ migratory neurons are abundant in the posterior cortical streams.**

**A.** Confocal images from the P0 piglet brain; coronal sections were immunostained for COUP-TFII, SP8, and DCX expression. The rostral migratory stream (RMS) and olfactory bulb (OB) glomeruli (delineated with a white dashed line) are populated by COUP-TFII-SP8+ DCX+ cells, as has been previously reported (*22*, *26*). COUP-TFII+SP8+ and COUP-TFII+SP8- DCX+ cells are rare in the dorsal and ventral tips extending from the olfactory ventricle. Scale bar, 50µm.

**B.** Quantification of subpopulations of DCX+ cells, differentiated by selected transcription factor (TF) expression, across anterior-posterior migratory streams in the P0 piglet brain. COUP-TFII-SP8+ DCX+ cells are more abundant in anterior migratory streams, like the RMS, whereas COUP-TFII+SP8+ DCX+ cells are found predominantly in posterior cortical streams.

**C.** Wide-field images of DCX+ cells expressing COUP-TFII and SP8 in the anterior and posterior cingulated cortex (CC), ventral streams, and temporal cortex 9TC) of the P0 piglet brain. Scale bar, 100µm.

**Figure S11.**

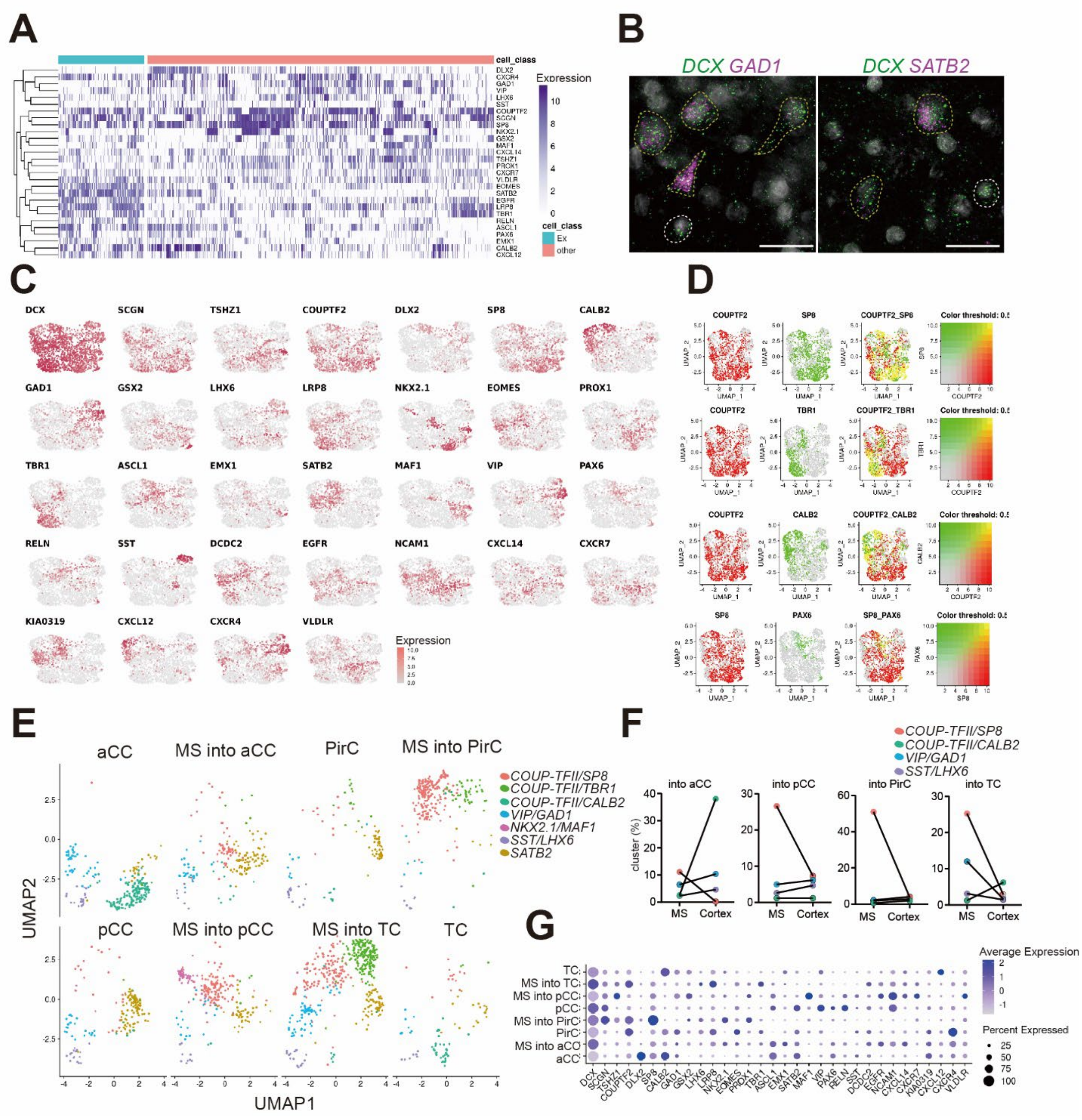


**Fig. S11. Spatial transcriptomics on P2 piglet brain uncovers heterogenous CGE-associated subpopulations across postnatal cortical streams.**

**A.** Heatmap depicting expression pattern of major cell type marker genes across the 1,992 *DCX*+ cells analyzed by Hi-Plex from P2 piglet brain. Migratory neurons cluster into *SATB2*-expressing excitatory (Ex; blue box) and GE marker-expressing inhibitory identities (red box).

**B.** Confocal images showing *DCX+GAD1+* migratory neurons (left; yellow dashed line) and *DCX+SATB2+* migratory excitatory neurons (right; yellow dashed line). Scale bar, 50µm.

**C.** Feature plots of 32 genes associated with interneuron versus excitatory neuron identity, and neuronal migration (see Table S5).

**D.** Feature plots illustrating degrees of gene co-expression. Yellow signal indicates that two genes are co-expressed. *COUP-TFII* is largely co-expressed with *SP8*, and only partially with *TBR1* and *CALB2*. The population co-expressing SP8 and PAX6, associated with dorsal lateral ganglionic eminence (dLGE), is small.
**E.** UMAP projection by different regions. Each migratory stream (MS) has an abundant *COUP-TFII/SP8* cluster. *COUP-TFII/TBR1* cluster is specified in the ventral MS, especially in the temporal cortex (TC). A small population of *NKX2.1/Maf1* is populated in the dorsal stream into the posterior cingulate cortex (pCC). Anterior CC (aCC); piriform cortex (PirC).
**F.** Quantification of cluster proportions in the migratory streams and their targeted cortical regions. The proportion of the *COUP-TFII/SP8* cluster gradually decreases along all migratory trajectories. Migratory stream (MS); anterior cingulate cortex (aCC); posterior CC (pCC); piriform cortex (PirC); temporal cortex (TC).
**G.** Dot plot illustrating the expression pattern of selected 32 genes across the different regions. Migratory stream (MS); anterior cingulate cortex (aCC); posterior CC (pCC); piriform cortex (PirC); temporal cortex (TC).

**Table S1. Comparative structural features across species.**

| | brain weight (g) | | gyrification index (GI) | | gray matter thickness (mm) | |
|---|---|---|---|---|---|---|
| Age | newborn | adult | newborn | adult | newborn | adult |
| Home Sapiens | 350-400 | 1300-1400 | 1.92±0.06 | 2.8 | 1.45±0.02 | 3.67±0.04 |
| Sus Scrofa | 32.5 | 180 | 1.86±0.03 | 2.16 | 1.34±0.02 | 1.56±0.02 |
| Macca Mulatta | 54 | 90-97 | 1.76±0.04 | 1.79 | 1.81±0.03 | 2.37±0.19 |
| Callithrix jacchus | 3.4 | 8 | 1.08±0.05 | 1.17 | 1.45±0.02 | 1.6 |
| Mus Musculus | 0.2 | 0.4 | 1.01±0.01 | 1.03 | 0.36±0.01 | 0.8-0.9 |

**Table S2. Comparative Arc features across species.**

| | Arc area ratio (%) | tier ratio in wedge area | | BV ratio in wedge area | MANOVA analysis |
|---|---|---|---|---|---|
| | | tier1 | tier2-3 | | |
| Home Sapiens | 0.63±0.04 | 0.13±0.01 | 0.87±0.01 | 0.025±0.007 | NA |
| Sus Scrofa | 0.44±0.12 | 0.22±0.02 | 0.77±0.02 | 0.011±0.001 | Pillai's, $F(3,4) = 3.98$, $p=0.1075$ |
| Callithrix jacchus | 0.18±0.08 | 0.97±0.01 | 0.02±0.01 | 0.003±0.001 | Pillai's, $F(3,6) = 398.82$, $p = 6.21x10\text{-}7$*** |
| Mus Musculus | 0.16±0.04 | 0.92±0.04 | 0.07±0.04 | 0.003±0.001 | Pillai's, $F(3,5) = 94.2$, $p = 8.07x10\text{-}5$*** |

**Table S3. Clinical and experimental demographics of collected human specimens**

| | Case No. | Age | Gender | Clinical History | Neuropathological diagnosis | Experimental Use |
|---|---|---|---|---|---|---|
| Human | 1 | 37GW+2 days | F | autopsy | Control | Nissl staining/TFs quantification/measurement of Arc area and BV ratio |
| | 2 | 0 day | F | left renal hypoplasia and right renal agenesis | Control | Nissl staining/RNAscope/TFs quantification/measurement of Arc area and BV ratio |
| | 3 | 37GW | F | Respiratory failure | Control | DCX mapping/TFs quantification/measurement of Arc area and BV ratio |
| | 4 | 36GW+2 weeks | F | cardiac anomaly | Control | Nissl staining/RNAscope/TFs quantification/measurement of Arc area and BV ratio |
| | 5 | 40 days | F | autopsy | Control | TF quantification |
| | 6 | 7 months | M | wiskott aldrich | Control | DCX intensity quantification |

**Table S3. Primary antibodies used in this study.**

| Primary Ab | Species | Dilution | Manufacturer | Cat.No. | Antigen Retrieval |
|---|---|---|---|---|---|
| ALDH1L1 | Mouse | 1:500 | Antibodies Inc | 75-140 | 10 min |
| a-SMA | Mouse | 1:500 | Sigma | a2547 | None |
| COUP-TFII | Mouse | 1:100 | R&D Systems | PP-H7147-00 | 11 min |
| BLBP | Rabbit | 1:500 | Abcam | ab32423 | None |
| Doublecortin | Rabbit | 1:200 | Cell Signaling | 4604 | None |
| Doublecortin | Guinea Pig | 1:200 | Millipore | ab2253 | None |
| DLX2 | Mouse | 1:200 | Santa Cruz | sc-393879 | 10 min |
| EMX1 | Mouse | 1:200 | Santa Cruz | sc-398115 | 10 min |
| FOXP2 | Rabbit | 1:500 | Millipore | ABE74 | 10 min |
| GFAP | Chicken | 1:750 | Abcam | ab4674 | 10 min |
| GSH2 | Rabbit | 1:200 | Abcam | ab26255 | 10 min |
| IBA1 | Goat | 1:250 | Abcam | ab5076 | 10 min |
| LHX6 | Mouse | 1:200 | Santa Cruz | sc-271433 | 10 min |
| MEIS2 | Mouse | 1:500 | Sigma | WH0004212M1 | 10 min |
| Mki67 | Rabbit | 1:200 | Abcam | ab15580 | 10 min |
| NeuN | Chicken | 1:500 | Millipore | ABN91 | None |
| NKX2.1 | Rabbit | 1:100 | Santa Cruz | SC-13040 | 11 min |
| Olig2 | Rabbit | 1:200 | Millipore | AB9610 | None |
| PAX6 | Mouse | 1:200 | Abcam | ab78545 | 10 min |
| PROX1 | goat | 1:200 | R&D | AF2727 | 10 min |
| PSA-NCAM | Mouse | 1:500 | Millipore | MAB5324 | None |
| SCGN | Mouse | 1:500 | Abcam | ab244219 | 10 min |
| SP8 | Goat | 1:100 | Santa Cruz | SC-104661 | 10 min |
| TSHZ1 | rabbit | 1:200 | Abcam | ab140196 | 10 min |

**Table S5. 32 gene lists for HiPlex RNAscope**

| Cycle 1 (plex 1-12) | Channel | Label Probe | Positive | Category |
|---|---|---|---|---|
| Round 1 | T1 | 488 | DCX | migratory neuron |
| | T2 | 550 | TSHZ1 | interneuron |
| | T3 | 647 | SCGN | interneuron |
| Round 2 | T4 | 488 | DLX2 | interneuron |
| | T5 | 550 | COUP-TFII | interneuron |
| | T6 | 647 | SP8 | interneuron |
| Round 3 | T7 | 488 | CALB2 | interneuron |
| | T8 | 550 | GAD1 | interneuron |
| | T9 | 647 | GSX2 | interneuron |
| Round 4 | T10 | 488 | LRP8 | migration-related receptor |
| | T11 | 550 | LHX6 | interneuron |
| | T12 | 647 | NKX2.1 | interneuron |

| Cycle 2 (plex 1-12) | Channel | Label Probe | Positive | Category |
|---|---|---|---|---|
| Round 1 | T1 | 488 | EOMES | excitatory neuron |
| | T2 | 550 | PROX1 | interneuron |
| | T3 | 647 | TBR1 | excitatory neuron |
| Round 2 | T4 | 488 | ASCL1 | interneuron |
| | T5 | 550 | SATB2 | excitatory neuron |
| | T6 | 647 | EMX1 | excitatory neuron |
| Round 3 | T7 | 488 | NA (negative ctl) | NA |
| | T8 | 550 | VIP | interneuron |
| | T9 | 647 | MAF1 | interneuron |
| Round 4 | T10 | 488 | PAX6 | interneuron |
| | T11 | 550 | RELN | migration-related chemokine |
| | T12 | 647 | SST | interneuron |

| Cycle 3 (plex 1-12) | Channel | Label Probe | Positive | Category |
|---|---|---|---|---|
| Round 1 | T1 | 488 | DCDC2 | migration |
| | T2 | 550 | NCAM1 | migration |
| | T3 | 647 | EGFR | migration |
| Round 2 | T4 | 488 | CXCL14 | migration-related chemokine |
| | T5 | 550 | KIA0319 | migration |
| | T6 | 647 | CXCR7 | migration-related receptor |
| Round 3 | T7 | 488 | CXCL12 | migration-related chemokine |
| | T8 | 550 | CXCR4 | migration-related receptor |
| | T9 | 647 | VLDLR | migration-related receptor |

**Movie S1.**
3D rendering of DCX+ cells (green) and BLBP+ cells (magenta) in dorsal streams from P0 piglet Arc.

**Movie S2.**
3D rendering of DCX+ cells (green) and BLBP+ cells (magenta) in ventral streams from P0 piglet Arc.

**Movie S3.**
Time-lapse imaging showing migrating neurons in P0 piglet organotypic slice culture. The area imaged is the dorsal side of the Arc. The cell-dense region is the Arc, and the white line delineates the boundary of the Arc. Note that labeled cells (•) are traveling in a dorsal direction, away from the Arc. The movie spans 34 hours.

**Movie S4.**
Time-lapse imaging showing migrating neurons in P0 piglet organotypic slice culture. The area imaged is the dorsal side of the Arc. The cell-dense region is the Arc, and the white line delineates the boundary of the Arc. Note that labeled cells (•) are traveling in a dorsal direction, away from the Arc. The movie spans 34 hours.

**Movie S5**
Light sheet imaging of a clarified P0 piglet brain from anterior to posterior. Black signal is DCX immunoreactivity. The stars (*) indicate dorsal streams of DCX+ cells from the Arc to the cingulate cortex and ventral streams of DCX+ cells from the Arc to the piriform cortex.
